# Single-nucleus transcriptomics reveals convergent effects of THC exposure and Reelin signaling on nucleus accumbens maturation in adolescence

**DOI:** 10.1101/2025.04.03.646846

**Authors:** Yanning Zuo, Avraham Libster, Arnav Gurha, Numaan Formoli, Leanne Liaw, Daimeng Sun, Andrew Turner, Attilio Iemolo, Francesca Telese

**Affiliations:** Department of Psychiatry, University of California, San Diego, La Jolla, California, USA

## Abstract

The nucleus accumbens undergoes extensive maturation during adolescence, but how drug exposure and genetic vulnerability interact to shape this process remains poorly understood. Here, we used single-nucleus RNA sequencing to examine the effects of chronic adolescent tetrahydrocannabinol (THC) exposure and reduced Reelin signaling in mice. THC produced broader transcriptional changes than Reelin haploinsufficiency, particularly in medium spiny neurons (MSNs). Analysis of cell-cell communication identified a THC-sensitive signaling program in which inhibitory interneurons were the principal receivers of MSN-derived signals related to axon-guidance and synaptic maturation. Despite the dominant effect of THC, both perturbations converged on shared gene networks linked to human genetic risk for substance use and psychiatric disorders. These effects were strongest in a population of immature neurons that we confirmed are generated in the adolescent nucleus accumbens and decline in adulthood. These findings show that adolescent THC exposure and Reelin signaling converge on transcriptional programs that regulate late neuronal maturation in striatal circuits.

## Introduction

Adolescence is a critical developmental period during which the brain undergoes extensive structural and functional remodeling to support adult cognitive and emotional behaviors(*1, 2*). The enhanced plasticity during this period increases vulnerability to environmental stressors, including exposure to substances of abuse. Epidemiological studies consistently associate frequent and heavy cannabis use in adolescence with elevated risk for neuropsychiatric disorders later in life, including psychosis and substance use disorders(*3–7*). Although these studies do not establish causality, converging evidence indicates that genetic factors modulate the relationship between cannabis exposure and long-term psychiatric outcomes(*8–10*).

Preclinical studies demonstrate that adolescent exposure to tetrahydrocannabinol (THC), the primary psychoactive component of cannabis, produces persistent impairments in cognitive and social behaviors(*11, 12*). THC exposure during adolescence alters brain structure and molecular signaling, including remodeling dendritic architecture in the prefrontal cortex and inducing long-lasting transcriptional changes across multiple brain regions(*11, 13–15*).

Among the brain regions sensitive to adolescent THC exposure(*11*), the nucleus accumbens (NAc) plays a central role due to its involvement in reward processing and motivated behaviors that are often dysregulated by substances of abuse(*16–18*). Alterations in gene regulatory programs within specific NAc cell types during adolescence may therefore have enduring consequences for behavioral regulation(*11, 19, 20*). Advances in single-nucleus RNA sequencing (snRNA-seq) have enabled high-resolution characterization of cellular diversity within the NAc, including molecular adaptations in the context of drug exposure, particularly within medium spiny neurons (MSNs) (*21–24*). However, how adolescent THC exposure shapes cell type-specific transcriptional programs in the NAc and how genetic risk modifies these responses remain largely unexplored.

Reelin represents a strong candidate modulator of these effects. This extracellular matrix protein regulates neuronal migration during development, modulates synaptic plasticity, and influences gene expression in both the developing and adult brain (*26–29*). Human genetic studies increasingly link variation in the RELN locus to risk for neuropsychiatric disorders(*30, 31*). In mice, haploinsufficiency of *Reln* alters behavioral responses to drugs of abuse, including psychostimulants and synthetic cannabinoids(*12, 25, 32–34*). These findings suggest that reduced Reelin signaling may amplify the developmental impact of adolescent cannabis exposure, providing a compelling framework for examining gene-environment interactions at the molecular level.

In this study, we utilized snRNA-seq to investigate how adolescent THC exposure alters transcriptional programs across NAc cell types in wild-type (WT) and *Reln* haploinsufficient (*Reln*+/−) mice. We performed differential gene expression, cell-cell communication and gene co-expression network analyses to identify cell type-specific molecular responses to THC exposure and reduced Reelin signaling. We then integrated these transcriptional signatures with human genome-wide association studies (GWAS) of substance use and psychiatric disorders. Finally, we perform histological validation of cell types identified in our study. Our findings provide novel mechanistic insights into cell type-specific responses underlying adolescent cannabis exposure and its relationship with the Reelin signaling.

## Results

### Single-nucleus RNA sequencing defines adolescent NAc cell types

We performed snRNA-seq on NAc tissue from mice treated chronically with THC (i.p., 10 mg/kg) or vehicle (**Fig. 1a**). Treatment began at postnatal day 30, corresponding to early adolescence in mice, and continued daily for 3 weeks. This exposure paradigm induces persistent behavioral and transcriptional changes, as shown previously(*11, 12*). We collected tissue one day after the final injection (postnatal day 51). The dataset comprised nuclei from 5 WT (n=2 vehicle, n=3 THC) and 8 *Reln* +/− (n=4 vehicle, n=4 THC) mice (**Data S1**). Following quality control filtering (**Fig. S1, Data S2**), the integrated dataset contained 61,911 high-quality nuclei clustered into transcriptionally distinct populations (**Fig. 1b-c, Fig. S2-4**). We annotated clusters using established marker genes (**Fig. 1d-e**). Most clusters belonged to the GABAergic neuronal lineage, defined by expression of *Slc32a1* and *Gad1* (**Fig. 1d**). MSNs accounted for more than 84% of neuronal nuclei and expressed lineage markers including *Foxp2, Ppp1r1b, and Bcl11b* (**Fig. 1d, Fig. S5a**). We further subdivided MSNs based on dopamine receptor expression, identifying five Drd1-expressing clusters (D1-MSN-1 to D1-MSN-5), three Drd2-expressing clusters (D2-MSN-1 to D2-MSN-3), and a smaller population of Drd3-expressing MSNs (D3-MSN, **Fig. 1d**). These subtypes formed transcriptional continua consistent with previous reports(*21, 22, 35, 36*), marked by *Tac1* enrichment in D1-MSNs and *Adora2* in D2-MSNs (**Fig. S5a)**. We also identified a distinct D1-MSN subpopulation with high *Grm8* expression (D1-MSN-5), consistent with prior reports(*37, 38*) (**Fig. S5a**). Additional subtype-specific markers, including *Ebf1, Onecut2, Sema5b, Greb1l,* and *Htr7*, further distinguished MSN populations **(Fig. S5a**).

**Fig. 1.**
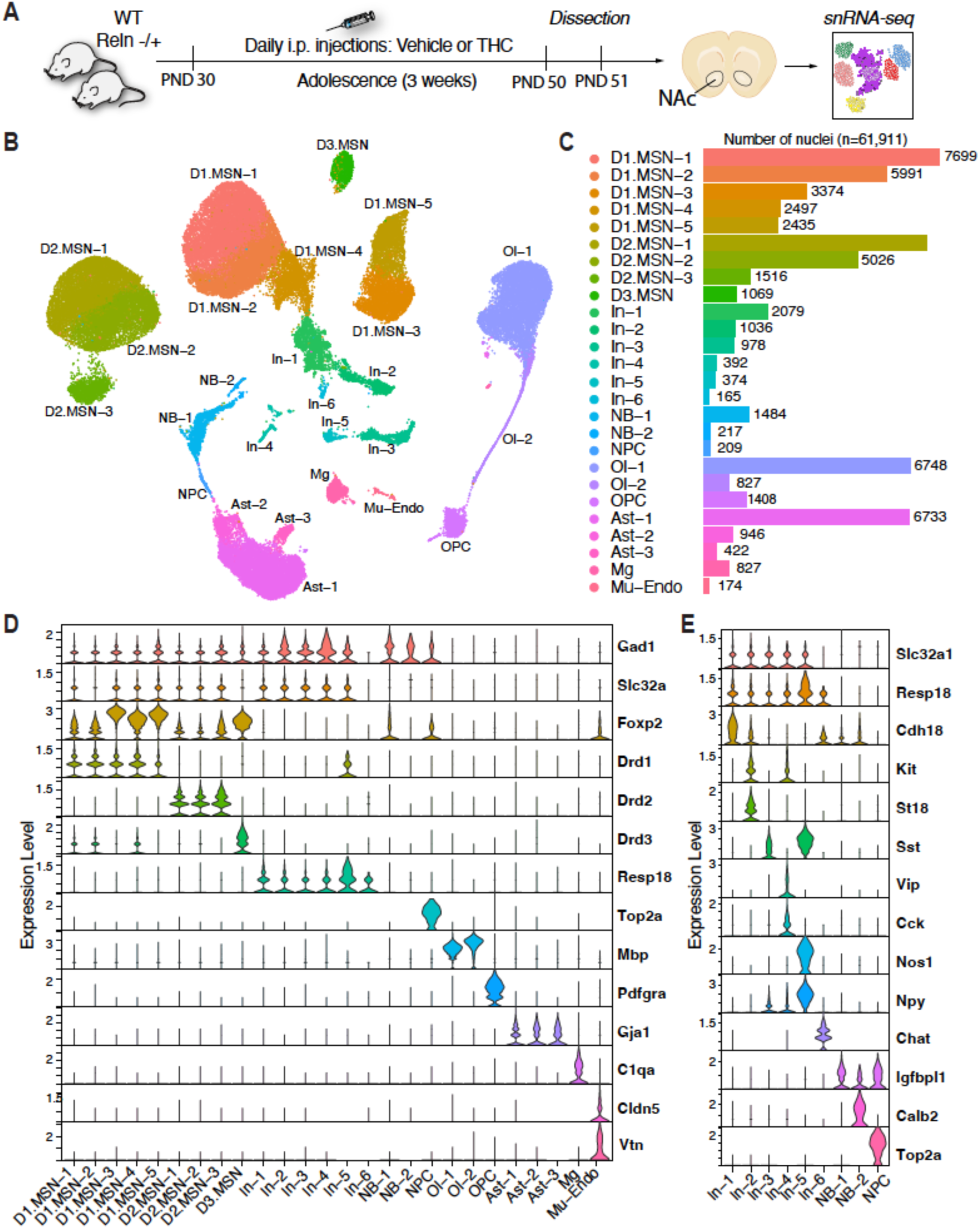
snRNA-seq profiling of adolescent NAc identifies diverse neuronal and non-neuronal cell types. **A)** Experimental timeline and workflow. THC or vehicle treatments were administered daily from PND30–50 (adolescence). NAc tissue was collected on PND51 and processed via snRNA-seq (10X Genomics). **B)** UMAP visualization of 61,911 integrated nuclei, colored by annotated cell clusters. **C)** Bar plot indicates the number of nuclei per identified cluster. **D)** Violin plots depicting expression levels of selected marker genes across annotated clusters, highlighting main cell type markers. Expression values shown as normalized RNA counts. **E)** Violin plots depicting expression levels of selected marker genes across interneurons (In), neuroblasts (NB) and neural progenitor cells (NPC). Expression values shown as normalized RNA counts.

Among non-neuronal cells, three astrocyte populations expressed *Gja1*, two oligodendrocyte populations expressed *Mbp*, and oligodendrocyte precursor cells (OPCs) were marked by *Pdgfra* expression (**Fig. 1d**). Additional cell types included *C1qa*-expressing microglia and mural-endothelial cells expressing *Cldn5* and *Vtn* (**Fig. 1d**).

Interneurons (In) represented a smaller neuronal fraction and expressed *Slc32a1* and *Resp18*(*21*). We identified six interneuron subtypes defined by distinct marker combinations: *Cdh18* (In-1)*, St18/Kit* (In-2), *Sst* (In-3), *Vip/Cck* (In-4), and *Nos1/Npy/Sst* (In-5), and a cholinergic population expressing *Chat* (In-6) (**Fig. 1e**). These subtypes correspond to previously described striatal interneuron classes and reflect functional diversity within the adolescent NAc(*39*).

We identified a population of proliferating cells expressing *Top2a,* consistent with neural progenitor cells (NPC), as reported previously in adolescent mouse midbrain regions(*40*) **(Fig. 1e**). We also detected two subsets of GABAergic neuroblasts (NB-1 and NB-2) expressing immature neuronal markers. NB-1 cells retained expression of the neural precursor marker *Igfbpl1*, whereas NB-2 expressed *Calb2*, consistent with early neuronal differentiation. These patterns suggest that the two neuroblast populations represent progressive stages of neuronal maturation.

*Reln* expression in the NAc was highly cell type-specific and localized primarily to subsets of D1-MSNs and interneurons (**Fig. S5b**), consistent with prior studies(*25, 32*). The Vip⁺/Cck⁺ interneuron cluster (In-4) showed robust co-expression of *Reln* and *Cnr1*, which encodes the cannabinoid receptor CB1. Analysis of adult human NAc snRNA-seq data(*41*) revealed a similar pattern, with RELN expression enriched in MSNs and co-expressed with CNR1 across several striatal neuronal classes (**Fig. S5c**). We validated these transcriptomic findings spatially using multiplexed RNAscope in mouse tissue. This analysis confirmed co-expression of *Reln* and *Cnr1* in both striatal and cortical regions (**Fig. S6-9**).

Comparative analysis with adult human NAc snRNA-seq datasets(*41*) demonstrated strong conservation of transcriptional profiles for D1- and D2-MSNs, interneurons, and glial populations (R² > 0.5, p < 0.05; **Fig. S10**). In contrast, Drd3-expressing MSNs and neuroblast populations showed limited conservation (R² < 0.3, p < 0.05, **Fig. S10**), likely reflecting differences in developmental stage and annotation resolution.

Together, these analyses define the cellular composition of the adolescent NAc and identify conserved neuronal subtypes, including *Reln/Cnr1*-expressing populations, relevant to adolescent cannabinoid exposure.

### THC and *Reln* haploinsufficiency alter NAc transcription

We performed differential gene expression analysis across all 26 annotated cell types using a Wilcoxon rank sum test (FDR < 0.05, |log₂FC| > 0.25; **Data S3-S4**). Comparison of THC- and vehicle-treated mice identified 2,231 differentially expressed genes (DEGs)-cell type pairs, corresponding to 1,493 unique genes (**Fig. S11)**. In contrast, comparison of *Reln*−/+ versus WT mice identified 626 DEGs-cell type pairs encompassing 506 unique genes (**Fig. S12**). Although THC exposure produced a broader transcriptional response than *Reln* haploinsufficiency, both perturbations preferentially affected MSN subtypes (**Fig. S13a**). A subset of 181 unique genes was differentially expressed in both contrasts (**Fig. S13b**). However, generalized linear modeling using DESeq2(*42*) detected no significant genotype × treatment interaction (data not shown).

Among the most strongly regulated THC-associated genes, the opioid neuropeptide gene *Penk* showed consistent upregulation across multiple MSN subtypes (**Fig. 2a, Fig. S11**). This pattern aligns with previous reports linking adolescent THC exposure to increased *Penk* expression in the NAc(*43*). In contrast, genotype-associated transcriptional changes included the expected downregulation of *Reln* and upregulation of *Cadm2* within MSNs (**Fig. 2b, Fig. S12)**. *Cadm2* encodes a synaptic cell adhesion molecule implicated by human genetic studies in lifetime cannabis use, risk-taking behaviors, and substance use vulnerability(*44, 45*).

**Fig. 2.**
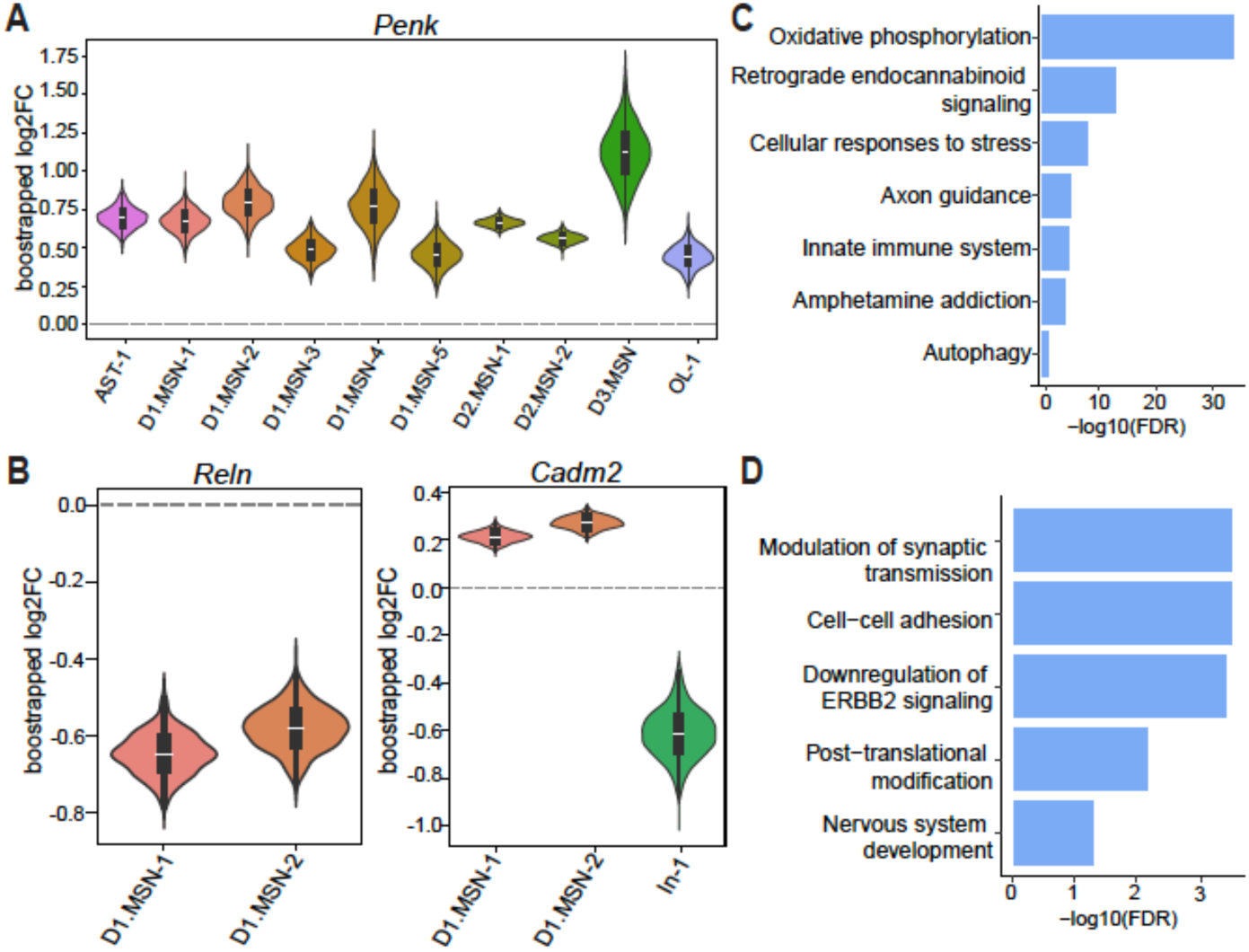
Differential gene expression analysis induced by adolescent THC exposure and Reln haploinsufficiency in the nucleus accumbens. (**A,B**) Violin plots with embedded boxplots showing the distribution of bootstrapped log₂ fold-change (log₂FC) estimates obtained using the Wilcoxon rank-sum test across 1,000 bootstrap iterations. Distributions are shown for cell types in which Penk (A) and Reln and Cadm2 (B) were significant (Q < 0.05) in at least 90% of bootstrap iterations. (**C**) Pathway enrichment analysis for THC-induced DEGs across neuronal cell types. (**D**) Pathway enrichment analysis for DEGs due to *Reln* haploinsufficiency. Bar plots in (C) and (D) show enriched pathways ranked by −log₁₀(FDR).

Pathway enrichment analysis revealed distinct molecular signatures associated with each perturbation. THC-regulated genes showed significant enrichment for oxidative phosphorylation, retrograde endocannabinoid signaling, axon guidance, and established addiction-related pathways (**Fig. 2c**). In contrast, genes associated with *Reln* haploinsufficiency were preferentially enriched for pathways related to synaptic transmission, cell-cell adhesion, and nervous system development (**Fig. 2d**).

Together, these results demonstrate that adolescent THC exposure induced more robust transcriptional changes than reduced Reelin signaling.

### THC alters interneuron-centered communication networks

To examine how adolescent THC exposure and *Reln* haploinsufficiency influence intercellular communication in the NAc, we integrated LIANA with Tensor-Cell2Cell, a context-aware tensor factorization framework that captures coordinated changes in ligand-receptor signaling across cell types and experimental conditions(*46, 47*). This analysis identified seven latent communication factors representing distinct patterns of cell-cell signaling (**Fig. S14**). No factors showed a significant association with genotype. In contrast, Factor 3 was selectively associated with THC exposure, exhibiting significantly higher receiver loadings in THC-treated mice compared with vehicle controls (adjusted *p* = 0.0118, **Fig. 3a**). Decomposition of Factor 3 revealed that interneurons served as the primary signal receivers, whereas D1- and D2- MSN subtypes acted as the principal signal senders (**Fig. 3b)**. Among interneurons, the In-4 subtype displayed the highest receiver loading. This population is characterized by co-expression of *Cnr1* and *Reln*, indicating preferential engagement of cannabinoid- and Reelin-associated signaling within this communication program. Functional annotation of the top 100 ligand-receptor-weighted genes contributing to Factor 3 revealed significant enrichment for pathways related to axon guidance, cell-cell adhesion, and calcium signaling. Protein-protein interaction network analysis further resolved six interconnected functional modules, including Ephrin receptor signaling, Neurexin-Neuroligin complexes, Ca²⁺/calmodulin signaling, axon guidance, retinoid metabolism, and Gα(i)-coupled receptor signaling (**Fig. 3d**). Intersection of the top 100 Factor 3-weighted genes with cell-type-resolved DEGs (FDR<5%) identified a significant overlap of 46 genes (**Fig. 3E**; **Data S5**). This overlap was significantly greater than expected by chance (Fisher’s exact test, odds ratio = 4.61, p = 1.0 × 10⁻¹²), indicating that the Factor 3 communication program is selectively engaged by THC-induced transcriptional changes. Disease enrichment analysis of these overlapping genes further linked this program to psychiatric and substance use-related disorders, including marijuana abuse (**Fig. 3F**).

**Fig. 3.**
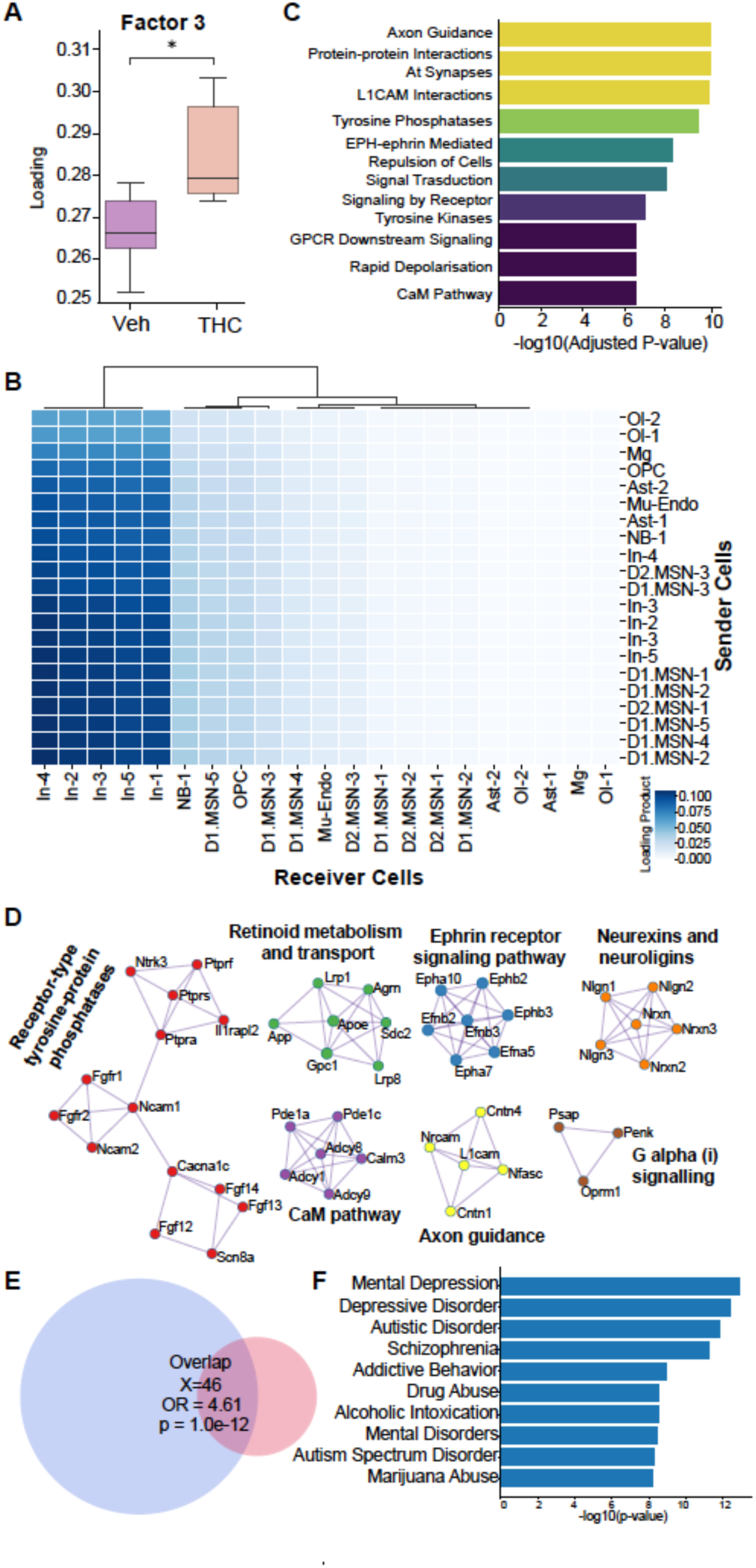
THC selectively alters an interneuron-centered cell-cell communication program. (**A**) Boxplots showing Factor 3 loadings derived from tensor-based cell-cell communication analysis, comparing vehicle- and THC-treated mice. Factor 3 loadings were significantly increased in the THC condition. (**B**) Heatmap of joint sender-receiver cell type loadings for Factor 3, representing a factor-specific network of intercellular communication, with interneurons showing the strongest receiver loadings and medium spiny neurons acting as predominant signal senders. (**C**) Overrepresentation analysis of the top 100 genes weighted by Factor 3 loadings, revealing enrichment for specific pathways. Bars indicate -log₁₀(adjusted p-value). (**D**) Protein-protein interaction networks generated from Factor 3-associated genes. Networks were clustered using the MCODE algorithm, highlighting interconnected functional modules. (**E**) Venn diagram showing the overlap between top 100 Factor 3-associated genes and DEGs following THC exposure. The overlap was significant. Odd ratio and p-value of the Fisher’s exact test are shown. (**F**) Disease enrichment analysis of the 46 DEGs using DisGeNET. Bars indicate −log₁₀(p-value).

Together, these analyses identify a THC-sensitive intercellular signaling program dominated by MSN-to-interneuron communication and centered on *Cnr1/Reln*-expressing interneurons.

### THC and *Reln* regulate cell type-specific gene networks

To assess whether THC-dependent intercellular effects reflect broader transcriptional coordination, we constructed gene co-expression networks using weighted gene co-expression network analysis (WGCNA) within each annotated cell type(*48*)(*49*) (**Fig. S15-21, Data S6**). Trait-module correlation analysis identified multiple gene modules significantly associated with THC treatment or *Reln* haploinsufficiency (**Fig. S22a-b, Data S7**), including three modules influenced by both perturbations (**Fig. 4a**). The NB-1_blue module showed opposing associations with treatment and genotype, exhibiting a negative correlation with THC exposure and a positive correlation with *Reln* haploinsufficiency (**Fig. 4a**). Genes within this module were enriched for pathways related to axon guidance, transcriptional regulation, and chromatin remodeling (**Fig. 4b**). In contrast, the D1-MSN-1_turquoise module showed the opposite pattern, with positive correlation to THC exposure and negative correlation with *Reln* genotype, and was enriched for pathways related to addiction, neural development, and synaptic transmission (**Fig. 4c**). The In-1_lightcyan module was negatively correlated with both perturbations but lacked clear functional enrichment and consisted primarily of non-coding RNAs.

**Fig. 4.**
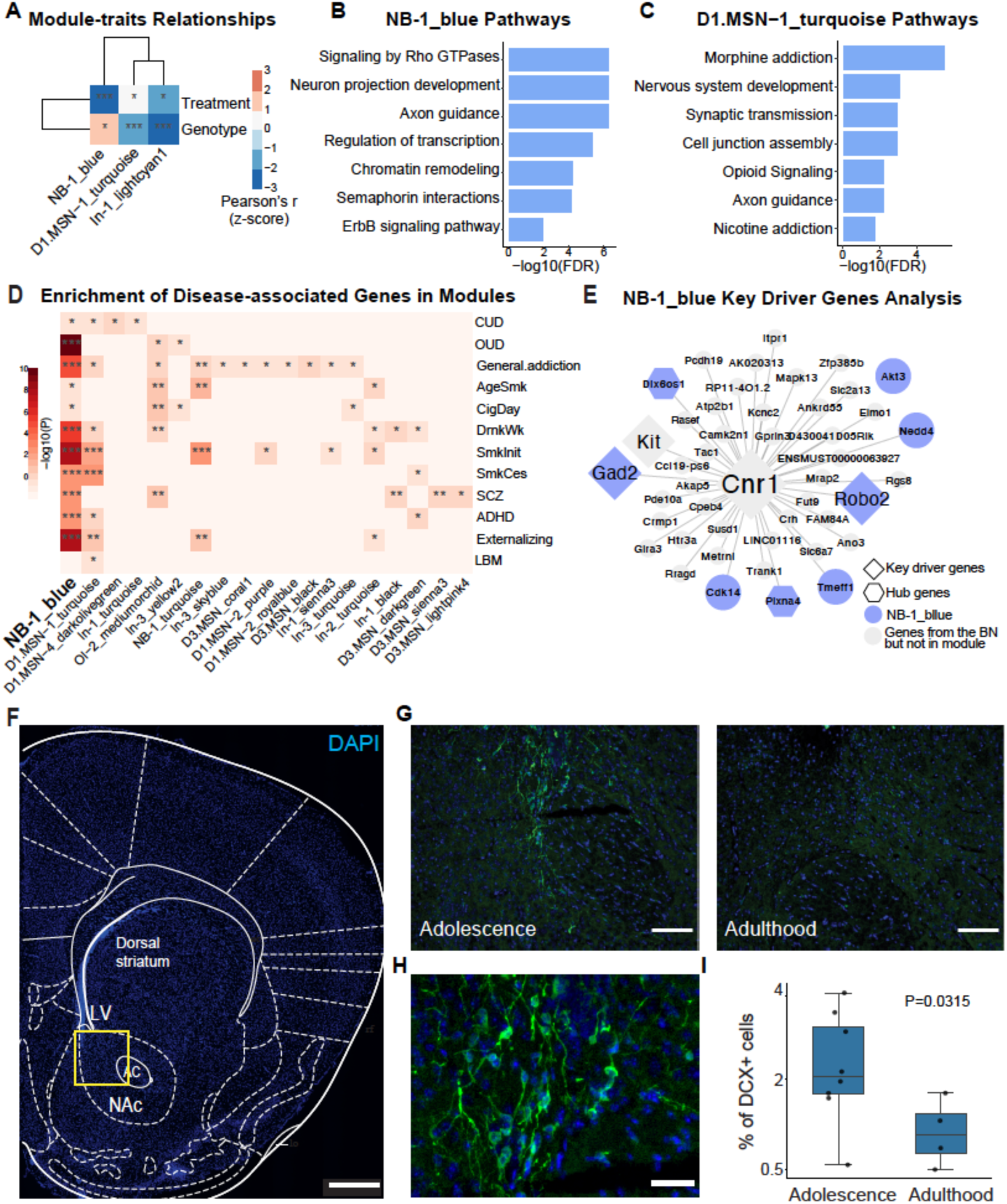
Cell type–specific gene co-expression networks link adolescent NAc maturation to genetic risk. (**A**) Heatmap showing module-trait correlations between WGCNA gene modules and experimental conditions (THC treatment and *Reln* haploinsufficiency). Colors represent Pearson’s correlation coefficients (z-scored); *p < 0.05, **p < 0.01, ***p < 0.001. (**B**) Pathway enrichment analysis of genes in the NB-1_blue module. (**C**) Pathway enrichment analysis of genes in the D1-MSN-1_turquoise module. Bar plots in (B) and (C) show enriched pathways ranked by −log₁₀(FDR). (**D**) Heatmap showing enrichment of WGCNA modules for disease-associated genes from genome-wide association studies using Mergeomics. Color indicates enrichment significance (−log₁₀(FDR)); Adjusted *p < 0.05, **p < 0.01, ***p < 0.001. (**E**) Bayesian network representation of the NB-1_blue module. Diamond nodes denote key driver genes; hexagons denote hub genes; blue nodes represent genes within the NB-1_blue module; gray nodes represent genes present in the Bayesian network but not assigned to these categories. (**F**) Representative coronal brain section showing DAPI staining (blue) with anatomical landmarks. LV, lateral ventricle; AC, anterior commissure; NAc, nucleus accumbens. Scale bar, 500 µm. (**G**) Representative images of DCX immunohistochemistry in the NAc during adolescence (left) and adulthood (right). Scale bar, 100 µm. (**H**) Higher-magnification image showing DCX-positive cells in the adolescent NAc. Scale bar, 20 µm. (**I**) Quantification of the percentage of DAPI-positive nuclei co-labeled with DCX in the NAc. Each point represents one mouse.

To evaluate the relevance of these transcriptional networks to human disease, we integrated treatment- and genotype-associated modules with genome-wide association study (GWAS) data using the Mergeomics framework(*50, 51*) (**Fig. S22c)**. Trait enrichment analysis across psychiatric and substance use GWAS datasets(*9, 52–57*) identified the NB-1_blue module as the most significantly associated (**Fig. 4d**), with strong enrichment for opioid use disorder, tobacco use and externalizing disorders (FDR < 0.05). This module was correlated with both THC exposure and *Reln* haploinsufficiency (**Fig. 4a**). Bayesian network analysis further identified *Cnr1* as a top key driver gene within the NB-1_blue module (**Fig. 4e**).

To validate the presence of immature neurons in the adolescent NAc, we performed immunohistochemistry (IHC) for doublecortin (DCX), a marker of migrating neuronal precursors(*58, 59*) (**Fig. 4f**). DCX-positive cells were present in the NAc and displayed elongated morphologies with leading processes (**Fig. 4g-h**), consistent with migratory phenotypes. Using CellProfiler(*60*), we quantified DCX-positive cells in adolescent (postnatal day 35–50) and adult (postnatal day 120) mice. The proportion of DCX-positive cells declined significantly from adolescence (2.4%) to adulthood (1.2%, p = 0.0315; **Fig. 4i**). To assess postnatal neurogenesis, we administered bromodeoxyuridine (BrdU) during the second week of adolescent THC exposure (**Fig. 5a**). We detected BrdU-positive cells ten days after the final BrdU injection and one day after the final THC or vehicle injection, indicating ongoing postnatal neurogenesis during adolescence. The number of BrdU+ and DCX+ cells did not differ between THC-treated and control groups (**Fig. 5b-c**).

**Fig. 5.**
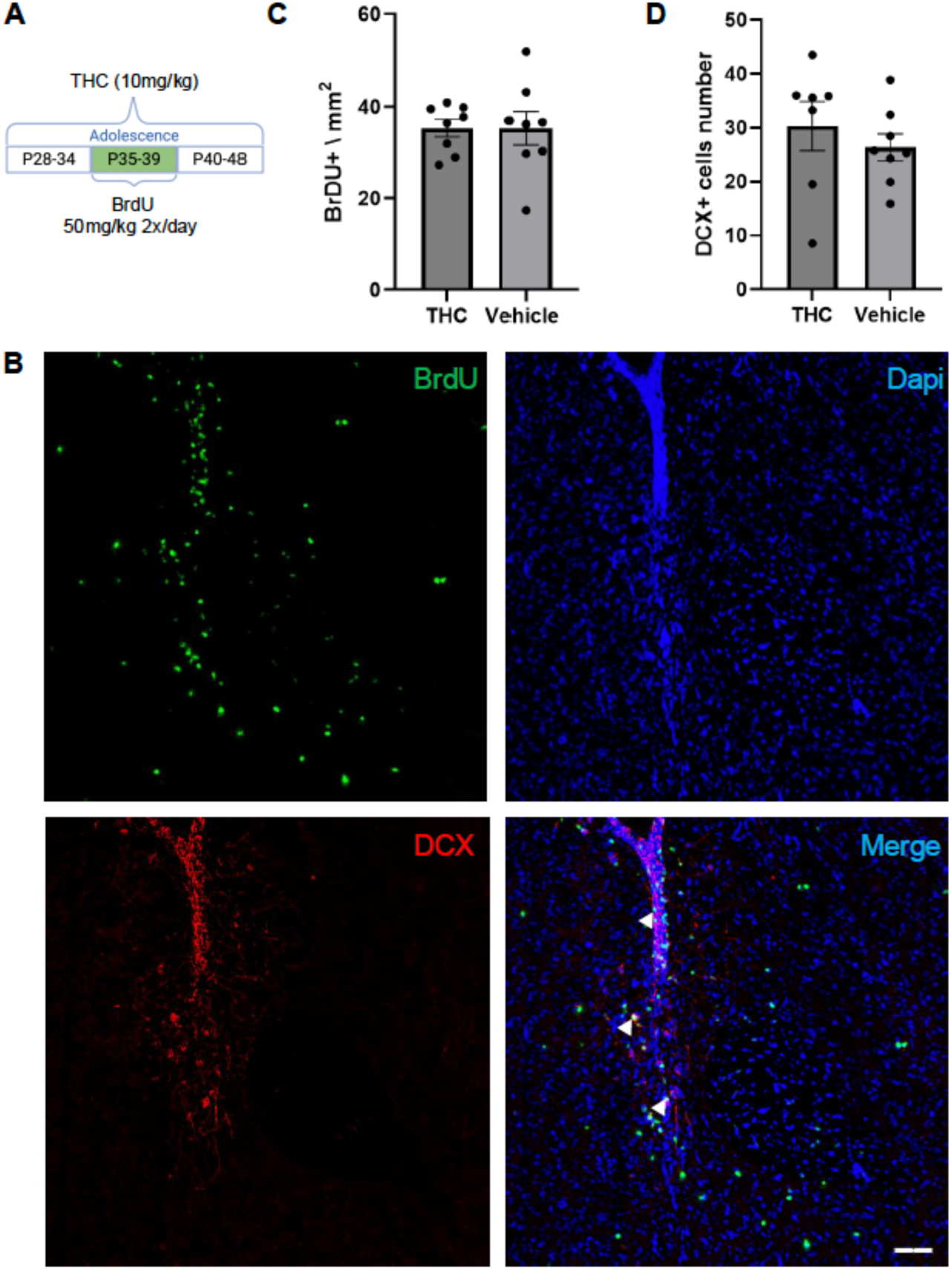
Adolescent THC exposure does not alter neuroblast proliferation or abundance in the NAc. (**A**) Experimental timeline illustrating adolescent THC exposure (10 mg/kg, i.p.) and BrdU administration (50 mg/kg, o.p., twice daily) during the second week of THC treatment (P35-39). Mice were perfused one day after the final THC injection (P49). (**B**) Representative images from the NAc showing BrdU (green), DCX (red), and DAPI (blue). Arrowheads indicate DCX-positive cells in proximity to BrdU-positive nuclei outside the ventricular wall zone. Scale bar, 50 µm. (**C**) Quantification of BrdU-positive cells per mm² in the NAc following THC or vehicle treatment. (**D**) Quantification of DCX-positive cells in the NAcfollowing THC or vehicle treatment. Each dot represents an individual mouse. Bars indicate mean ± SEM.

Together, these results show that gene networks associated with immature neuronal populations are sensitive to both THC exposure and reduced Reelin signaling and are linked to human genetic risk for psychiatric and substance use disorders.

## Discussion

Adolescence represents a prolonged window of NAc maturation during which environmental exposures can interact with genetic vulnerability to shape long-term circuit organization. Our findings indicate that adolescent THC exposure and reduced Reelin signaling reshapes developmental programs in the NAc by altering coordinated intercellular signaling and transcriptional networks.

A central insight from this work is that adolescent THC exposure preferentially perturbs intercellular communication programs, rather than acting solely through cell-intrinsic transcriptional changes. Tensor-based analysis identified a THC-sensitive signaling factor (Factor 3) associated with developmental pathways, such as axon guidance and synaptic maturation, and dominated by MSN-to-interneuron communication, with *Cnr1/Reln-*expressing interneurons emerging as the primary signal receivers. This result suggests that adolescent THC exposure alters how inhibitory interneurons integrate neuromodulatory inputs within developing NAc circuits.

Immature GABAergic neuroblasts also emerged as a population sensitive to both THC exposure and *Reln* haploinsufficiency. Multiple lines of evidence, including co-expression network analysis, enrichment for psychiatric and substance use GWAS signals, and histological validation using DCX and BrdU, support the interpretation that these cells represent a developmentally relevant population contributing to postnatal NAc maturation. Because neuroblasts did not express *Cnr1*, THC-associated transcriptional changes in this population likely arise indirectly through altered intercellular signaling. Importantly, THC exposure did not change neuroblast abundance, but instead modulated gene programs linked to axon guidance, synaptic adhesion, and chromatin regulation, consistent with altered maturation trajectories rather than altered neurogenic output.

Our data further clarify how THC exposure intersects with Reelin signaling. Although *Reln* haploinsufficiency alone did not strongly alter cell-cell communication factors, it biased the same developmental axis targeted by THC. Modules sensitive to both perturbations showed enrichment for human genetic risk associated with substance use and psychiatric disorders, linking experimental manipulations to clinically relevant pathways. These findings support a model in which reduced Reelin signaling alters the baseline developmental landscape, increasing susceptibility to environmental perturbation during adolescence.

Several genes linking intercellular signaling to transcriptional regulation showed strong sensitivity to THC exposure, including *Penk*, *Nrxn3*, and components of Eph-ephrin signaling pathways. Increased *Penk* expression in MSNs aligns with prior evidence that adolescent THC exposure enhances striatal opioid signaling and increases vulnerability to opioid self-administration(*43, 61*). In parallel, changes in *Nrxn3* and Eph-ephrin pathway genes point to disruption of synaptic adhesion and activity-dependent circuit refinement, processes essential for the stabilization of mature striatal connectivity(*62–64*). Notably, *NRXN3* genetic variants are strongly associated with substance use disorder and impulsivity (*65–71*). Together, these molecular changes suggest that adolescent THC exposure perturbs both neuromodulatory balance and structural synaptic programs within the NAc.

Several limitations warrant consideration. Although our analyses identify THC- and Reelin-sensitive communication and transcriptional programs, future studies using electrophysiology, circuit tracing, and spatial transcriptomics will be necessary to define how these molecular changes translate into functional connectivity. In addition, experimental validation of predicted ligand-receptor interactions will be required to establish causal mechanisms linking cannabinoid signaling, Reelin-responsive cells, and neuroblast maturation. Finally, while we focused on a developmentally relevant THC exposure paradigm, future work will need to address dose dependence, sex-specific effects, and the influence of different exposure windows.

In conclusion, our findings identify a cannabinoid-sensitive developmental signaling in the adolescent NAc that links *Cnr1/Reln-*expressing interneurons, neuroblast maturation, and addiction-related genetic pathways. By highlighting coordinated changes in intercellular communication and transcriptional maturation, our findings reveal how environmental exposures during adolescence interact with genetic vulnerability to increase long-term risk for substance use and psychiatric disorders.

## Materials and Methods

### Animals

All experimental procedures were approved by the institutional animal care and use committee at the University of California, San Diego. Mice were housed (3-4 per cage) under a 12h light/12h dark cycle and provided with food and water ad libitum.

For snRNA-seq experiments, the heterozygous reeler mouse(*29*) (*Reln* −/+) were bred in house using the B6C3Fe a/a-Relnrl/J line (The Jackson Laboratory, #000235). For snRNA-seq, we used 5 WT mice (n= 2 vehicle, n=3 THC), and 8 *Reln* −/+ mice (n=4 vehicle, n=4 THC). For DCX-IHC experiments in adolescent and adult mice we used 12 C57BL/6J male, and for BrdU experiments, we used 16 C57BL/6J male mice, purchased from Jackson Laboratory (#000664).

### Drug treatment protocol and experimental design

THC was supplied by the U.S. National Institute on Drug Abuse and prepared in a vehicle solution containing ethanol, tween, and 0.9% saline in a ratio of 1:1:18. The solution was freshly prepared on the day of administration. Specifically, we administered vehicle or THC (10 mg/kg) daily to mice by intraperitoneal injections in the early adolescent period, from PND 30 to PND 50 +/− 3 days, as previously reported(*11, 12, 72*). In our previous studies(*11, 12*), we employed the same drug treatment protocol and demonstrated that it resulted in long term behavioral deficits and plasma THC concentrations of 5–6 ng/ml one day after the final injection. While data on plasma THC levels in adolescent users remain limited, these findings align with studies on frequent adult cannabis users, which reported median plasma THC levels of ≥5.0 ng/ml for up to three days of abstinence(*73–75*). Despite the variability in THC content across cannabis products used by humans, the dosage used in this study is likely reflective of heavy cannabis use. In this study, we collected the brain tissues 1 day after the last injection.

### Single-nucleus RNA-seq data generation

NAc tissue was microdissected from coronal brain sections at approximately +1.78 mm from bregma and processed for snRNA-seq. Tissue punches from individual mice were homogenized and nuclei were isolated and permeabilized. Approximately 12,000 nuclei per sample were loaded onto a Chromium Controller, and snRNA-seq libraries were generated by the UC San Diego Center for Epigenomics using the 10x Genomics Chromium Single-Cell 3′ v3 chemistry following manufacturer protocols. Libraries were quality controlled for concentration and fragment size and sequenced on Illumina NextSeq 500 and NovaSeq 6000 platforms using paired-end reads (28 bp Read 1, 8 bp index, 91 bp Read 2).

### Bioinformatic analysis of snRNA-seq data

#### Raw data processing and quality control

We processed raw sequencing data using Cell Ranger (v6.0.2) and aligned reads to the mm10-3.0.0 reference genome. We integrated gene expression matrices from 13 samples using the aggr function in Cell Ranger (v6.0.1), yielding a combined dataset of 82,251 nuclei. We imported the merged gene expression matrix into Seurat (v4.9.9.9058) using Read10X_h5() and CreateSeuratObject(). We retained nuclei with 1,000-16,000 RNA counts, 500-6,000 detected genes, <2% mitochondrial gene content, <1% ribosomal small subunit (RPS) genes, and <1% ribosomal large subunit (RPL) genes. We detected potential doublets using DoubletFinder(*76*) (v2.0.3). We performed dimensionality reduction on each sample using the first 20 principal components. We set the artificial doublet rate (pN) to 0.25 and determined the optimal neighborhood size (pK) using paramSweep_v3(), summarizeSweep(), and find.pK(). Assuming a 5% doublet rate, we retained 68,917 nuclei after doublet removal.

#### snRNA-seq cell clustering, and cell annotation

We normalized data using SCTransform() with regularized negative binomial regression, regressing out mitochondrial gene percentage, sequencing depth, and ribosomal gene content. We performed principal component analysis using RunPCA() and used the first 50 PCs for graph-based clustering and visualization with FindNeighbors(), FindClusters(), and RunUMAP(). We used a clustering resolution of 0.7. Markers for each nuclear cluster were identified using ‘FindMarkers()’ in Seurat. We used a list of manually curated canonical markers to facilitate cell type annotation: Neuron: *Snap25*, *Syt1*; Glutamatergic excitatory neuron: *Slc17a7*; GABAergic interneuron: *Resp18*, *Gad1*; MSN: *Ppp1r1b, Foxp2, Bcl11b*; Drd1+ MSN: *Drd1, Pdyn, Tac1*; Drd2 MSN: *Drd2, Adora2a, Penk*; Oligodendrocyte: *Mbp*; OPC: *Pdgfra*; Astrocyte: *Gja1*; NPC *Top2a*; NB: *Igfbpl1*. Microglia: *C1qa*; Mural cell: *Vtn*; Endothelial cell: *Cldn5*. We excluded two clusters with high mitochondrial gene content and mixed neuronal-oligodendrocyte markers. We also excluded glutamatergic neuron clusters, as the NAc lacks intrinsic excitatory neurons. A total of 61,911 nuclei were retained for downstream analyses.

#### Differential gene expression analysis

We identified differentially expressed genes using FindMarkers() in Seurat for treatment (THC vs. vehicle) and genotype (*Reln⁺/−* vs. WT) within each cell type. We defined significance as |avg_log₂FC| > 0.25 with a false discovery rate (FDR) < 0.05. For visualization of DEGs, a bootstrapping analysis was performed within each cell type cluster identified in the dataset. This approach generates a distribution of gene expression statistics robust to single-cell sampling variability. For either treatment or genotype effect, bootstrapping was performed by sampling 61,911 nuclei with replacement for 1,000 iterations. In each iteration nuclei were independently sampled from within the constituent groups of comparison (THC vs Vehicle or WT vs HET), ensuring the original total number and proportion of nuclei in each group was maintained across all resamples. Differential gene analysis was performed using the FindMarkers function in Seurat (v5.3.0). The p values were calculated using the default non parametric Wilcoxon rank sum test. To prioritize the highly confident DEGs (FDR<0.05) we set a statistically significant threshold of a minimum of 900/1000 iterations (>90% frequency).

#### Weighted gene co-expression network analysis

We used the WGCN R package(*48*), as previously adapted for snRNA-seq(*49*). We normalized raw counts using NormalizeData() and log-transformed expression values. For each cell subtype, we selected nuclei within the 5th–40th percentile of RNA counts to exclude low-quality nuclei and potential doublets. We generated adjacency matrices using biweight midcorrelation and cell type-specific soft-thresholding powers. We calculated topological overlap matrices and identified gene modules using dynamic tree cutting (cutreeDynamic, method = hybrid, minModuleSize = 30, pamStage = F, pamRespectsDendro = T, deepSplit = 2, and cutHeight = 0.9999). We merged modules with eigengene correlations >0.85. We assessed module-trait associations using Pearson correlation between module eigengenes, THC treatment, and *Reln* genotype. We calculated z-scores of correlations across modules for visualization.

#### Pathway enrichment analysis of DEGs and WGCNA modules

We performed pathway enrichment analysis using the enrichR package(*77*). We queried KEGG_2021_Human, GO_Biological_Process_2021, BioCarta_2016, and Reactome_2022 databases. We adjusted p-values using the Benjamini–Hochberg method and considered adjusted p-values < 0.05 significant.

#### Integration with human GWAS data with Mergeomics

We integrated coexpression modules with human GWAS summary statistics using Mergeomics(*50*) R package. We analyzed GWAS datasets for cannabis use disorder (*9*), opioid use disorder (*54*), externalizing traits(*55*), general addiction risk factor(*57*), schizophrenia(*53*), ADHD(*56*), tobacco and alcohol use(*52*), and lean body mass(*78*). We pruned SNPs for linkage disequilibrium (r² < 0.5) and mapped SNPs to genes based on proximity (±20 kb) and nucleus accumbens–specific eQTLs and sQTLs from GTEx(*79*) v8. We assessed enrichment using Marker Set Enrichment Analysis (MSEA). We considered modules with FDR < 0.05 significantly enriched. We identified key driver genes using weighted Key Driver Analysis (wKDA) in Mergeomics with a brain-specific Bayesian gene network previously used in our study(*11*). We visualized networks using Cytoscape (v3.8.2).

#### Cross-species conservation analysis

We calculated cell type-specific t-statistics on SCTransformed expression values following established methods(*80*): We compared adolescent mouse nucleus accumbens cell types with adult human(*41*) NAc dataset(*40*). We used the cell type identities reported in the original study and applied SCTransform normalization, regressing out sequencing depth, donor, sex, processing batch, and ribosomal gene content (percent.RPL and percent.RPS). We calculated Pearson correlations using the top 100 upregulated genes per cell type.

#### Cell-cell communication analysis

We processed data in Python (v3.10.18) using Scanpy(*81*), LIANA+(*47*), and Tensor Cell2Cell(*46*). We retained cell types present in at least 70% of samples. We constructed ligand–receptor tensors from LIANA magnitude rank scores and identified seven communication factors using Tensor Cell2Cell. We assessed THC effects on factor loadings using t-tests with Benjamini-Hochberg correction. To characterize the biological pathways represented by Factor 3, we performed pathway overrepresentation Analysis (ORA) for Factor 3 using the Reactome 2022 database in EnrichR(*77*). Gene weights from Factor 3 loadings were parsed from ligand-receptor pairs, and unique genes were ranked by their maximum Factor 3 score. The top 100 genes were selected as the input “target list,” while the background consisted of all genes represented in the analysis. We performed protein-protein interaction analysis using Metascape(*82*). The top 100 genes ranked by Factor 3 loadings were intersected with DEGs (n=1,813, FDR<0.05) across cell types, and overlap significance was assessed using Fisher’s exact test using a background of 11,435 expressed genes detected across cell types. We performed disease enrichment analysis for the 46 overlapping genes between top 10 Factor 3 and DEGs using the Disgenet database in EnrichR(*77*).

### DCX Immunohistochemistry in adolescent and adult mice

We perfused 4 adolescent and 4 adult mice with 4% paraformaldehyde. Brains were post-fixed in 4% PFA, cryoprotected in sucrose, and sectioned coronally at 16 µm thickness. We stained sections with anti-DCX antibodies (Cell Signaling, 4604S). We acquired images using a Keyence BZ-X810 microscope. We quantified the DCX-positive cells using a customized CellProfiler (v4.2.5) pipeline(*60*). Immunofluorescence images were converted to grayscale and subjected to background correction to reduce illumination artifacts. Nuclei were identified from DAPI images using watershed-based segmentation and filtered by size to exclude debris and merged objects. DCX-positive cells were detected from fluorescence images using intensity-based thresholding and size filtering.

### BrdU labeling and immunohistochemistry

To label proliferating cells during adolescence, mice received intraperitoneal injections of BrdU (Sigma, B5002) at a dose of 50 mg/kg, administered twice daily for five consecutive days during the second week of the THC exposure period. Mice were perfused with 4% PFA one day after the final THC or vehicle injection. For each mouse, three sections spanning the nucleus accumbens were selected. For each section, both the left and right hemispheres were imaged, yielding up to six images per mouse. Images with tissue damage or poor staining quality were excluded from analysis. Immunohistochemistry was performed using antibodies against DCX (Cell Signaling Technology, 4604S) and BrdU (Sigma, 11170376001), with DAPI used for nuclear counterstaining, following previously reported protocols(*83*). Images were acquired using a Keyence BZ-X810 fluorescence microscope with a 20× objective, collecting stitched Z-stack images to capture the full section thickness. DCX-positive cells were quantified manually using the Cell Counter plugin in ImageJ. For DCX analysis, 8 vehicle-treated mice and 7 THC-treated mice were included; one THC-treated mouse was excluded due to insufficient staining quality. We counted only DCX-positive cells located within the nucleus accumbens parenchyma and excluded cells located within or immediately adjacent to the ventricular wall. The ventricular boundary was defined based on established anatomical morphology and local DAPI density within the same image. Cell counts were aggregated across sections for each animal and analyzed using GraphPad Prism.

### Statistical analysis

We performed statistical analyses in GraphPad Prism. We used unpaired two-tailed t-tests for DCX or BrdU quantification. We defined p ≤ 0.05 as statistically significant.

## Acknowledgments

We thank Havilah Taylor and Patricia Montilla Perez for technical assistance with mouse colony management. We acknowledge support by the UCSD Center for Epigenomics for generating snRNA-seq libraries and the UCSD IGM Genomics Center for sequencing the snRNA-seq libraries.

## Author contributions

Conceptualization: F.T, Y.Z, A.L Methodology: Y.Z, A.L, AG, F.T. Investigation: Y.Z, A.L, N.F, A.G, L.L, D.S, A.T, A.I. Formal analysis: Y.Z, A.L., A.G., D.S., F.T. Visualization: Y.Z, A.L., A.G., F.T. Supervision: Y.Z., A.L., F.T. Writing-original draft: FT. Writing-review and editing: Y.Z., F.T.

## Data and materials availability

All data needed to evaluate the conclusions in the paper are present in the paper and/or the Supplementary Materials. The raw and processed snRNA-seq data along with related sample information in this study is deposited in the Gene Expression Omnibus (GEO: GSE244913) database. **Reviewer access token: sriviymatjarjen**

