## supplementary materials for "Single-nucleus transcriptomics reveals convergent effects of THC exposure and Reelin signaling on nucleus accumbens maturation in adolescence"

Yanning Zuo, *et al.*

**This PDF file includes:**

Figs. S1 to S22

**Other Supplementary Materials for this manuscript include the following:**

Data S1 to S7

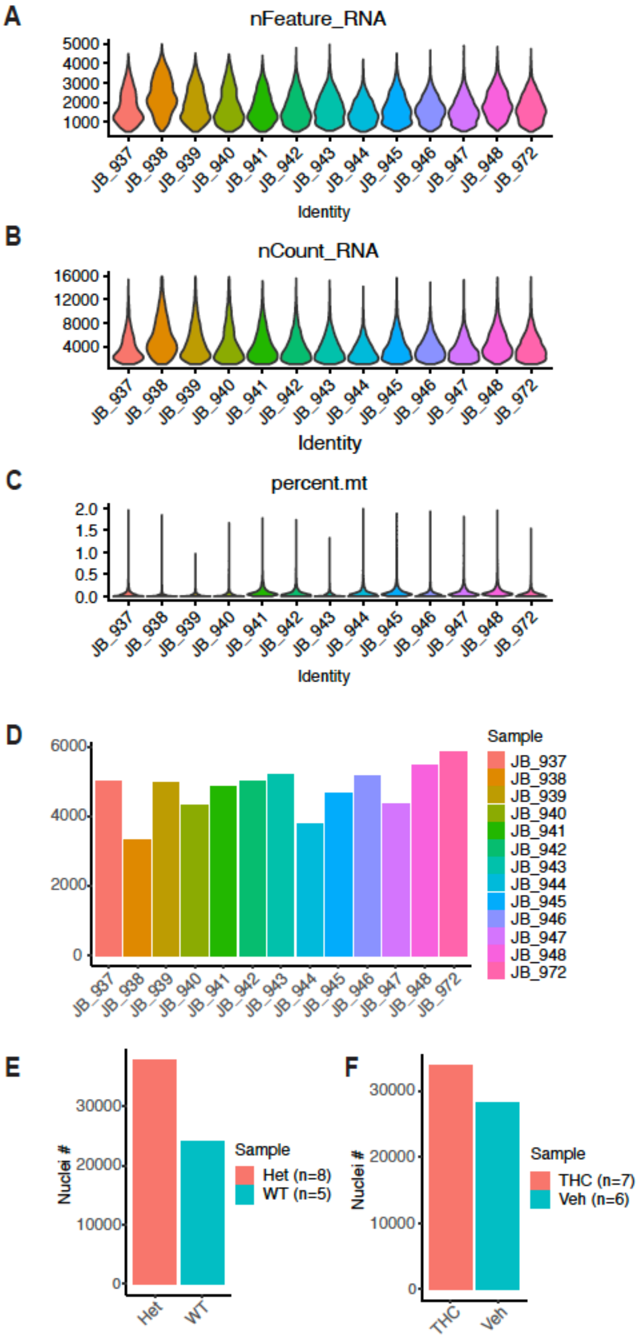


**Fig. S1. Quality control metrics for snRNA-seq datasets.** (**A–C**) Violin plots showing the distribution of detected genes per nucleus (nFeature_RNA), total RNA counts (nCount_RNA), and mitochondrial gene percentage (percent.mt) for each individual sample. (**D**) Total number of high-quality nuclei retained per sample after quality control filtering. (**E**) Total nuclei counts aggregated by genotype (Reln$^{+}$/−, n = 8; WT, n = 5). (**F**) Total nuclei count aggregated by treatment condition (THC, n = 7; vehicle, n = 6).

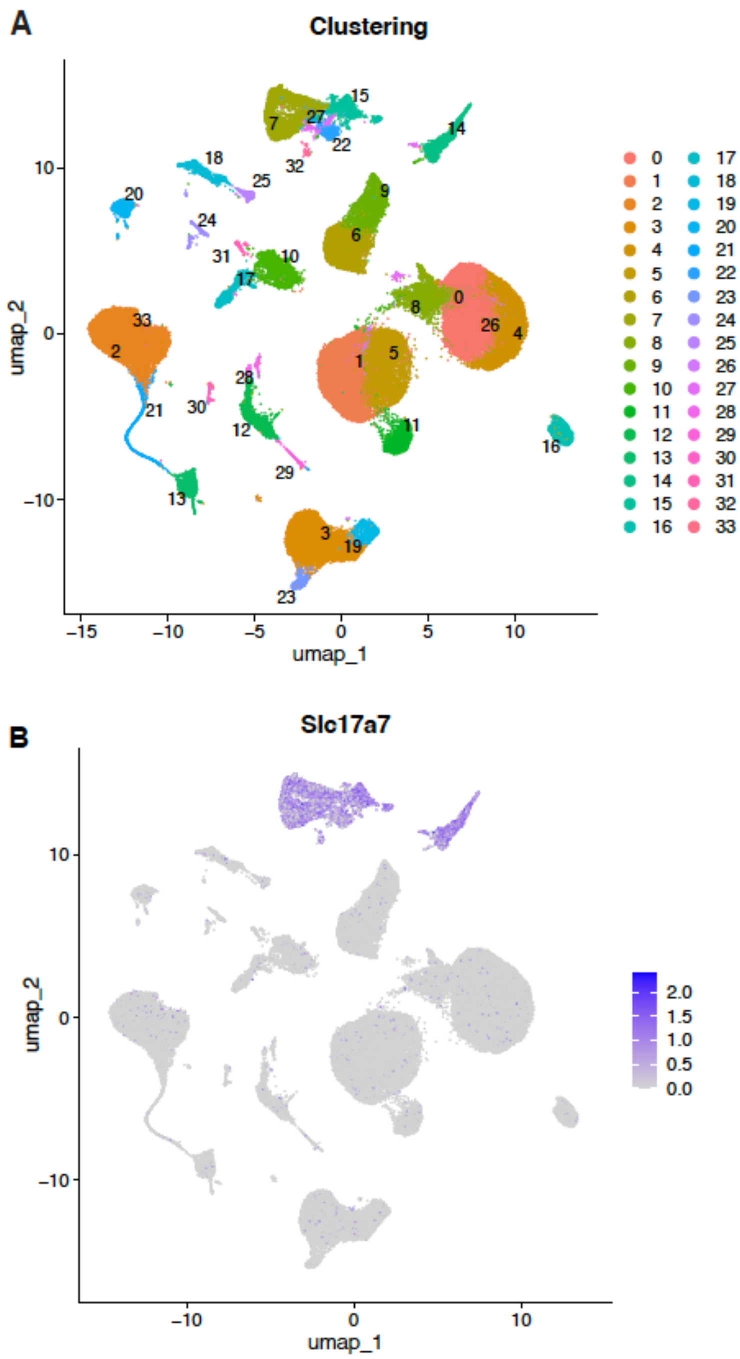


**Fig. S2. Clustering resolution and identification of glutamatergic populations.** (**A**) UMAP visualization showing high-resolution clustering of snRNA-seq data from the adolescent nucleus accumbens. Each cluster is labeled by numeric identifier. (**B**) Feature plot showing expression of *Slc17a7*, a marker of glutamatergic neurons, highlighting a small subset of cells with excitatory neuron signatures.

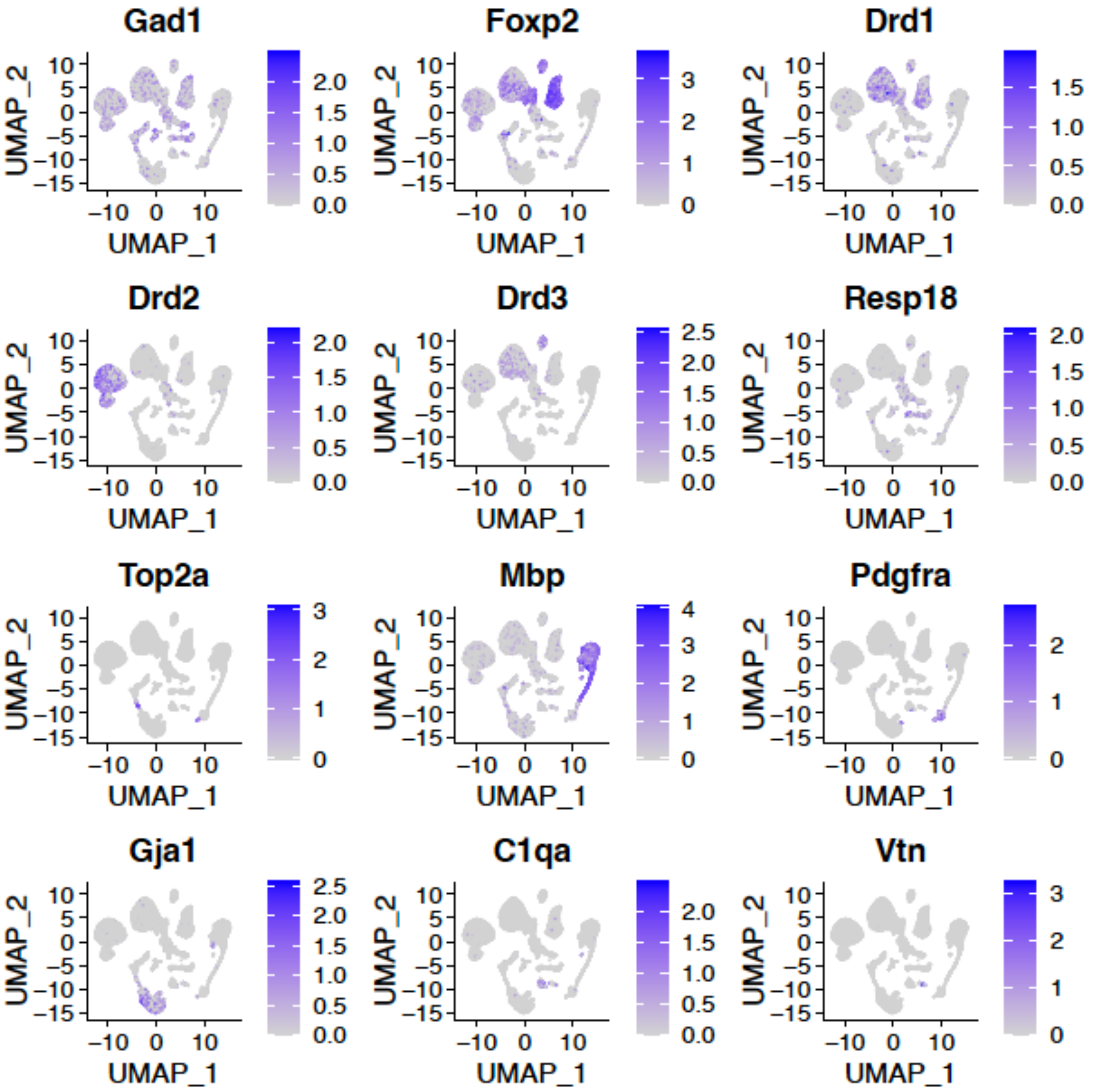


**Fig. S3. Marker gene expression used for cell type annotation in the adolescent nucleus accumbens.** Feature plots showing UMAP visualizations of expression for representative cekk type marker genes used to annotate cell clusters, including *Gad1, Foxp2, Drd1, Drd2, Drd3*, and *Resp18* (neuronal populations); *Top2a* (neural progenitors); *Mbp* and *Pdgfra* (oligodendrocytes and oligodendrocyte precursor cells); *Gja1* (astrocytes); *C1qa* (microglia); and *Vtn* (mural/endothelial cells).

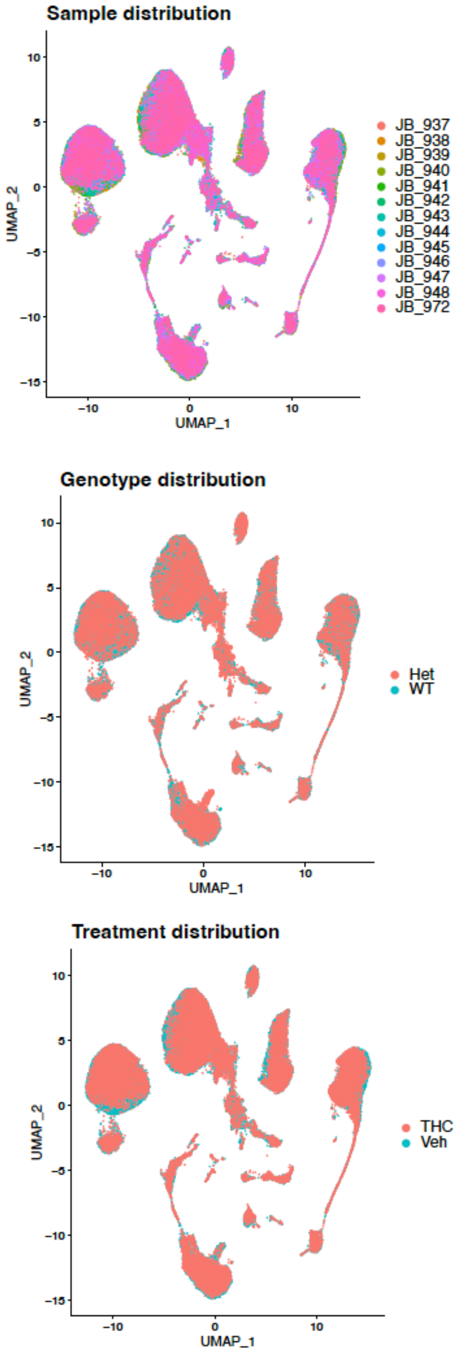


**Fig. S4. Integration and distribution of nuclei across samples, genotype, and treatment**. UMAP visualizations showing the distribution of nuclei colored by individual sample (top), genotype (middle), and treatment condition (bottom). Plots demonstrate consistent representation and integration of nuclei across clusters, with no evident segregation by sample, genotype, or treatment.

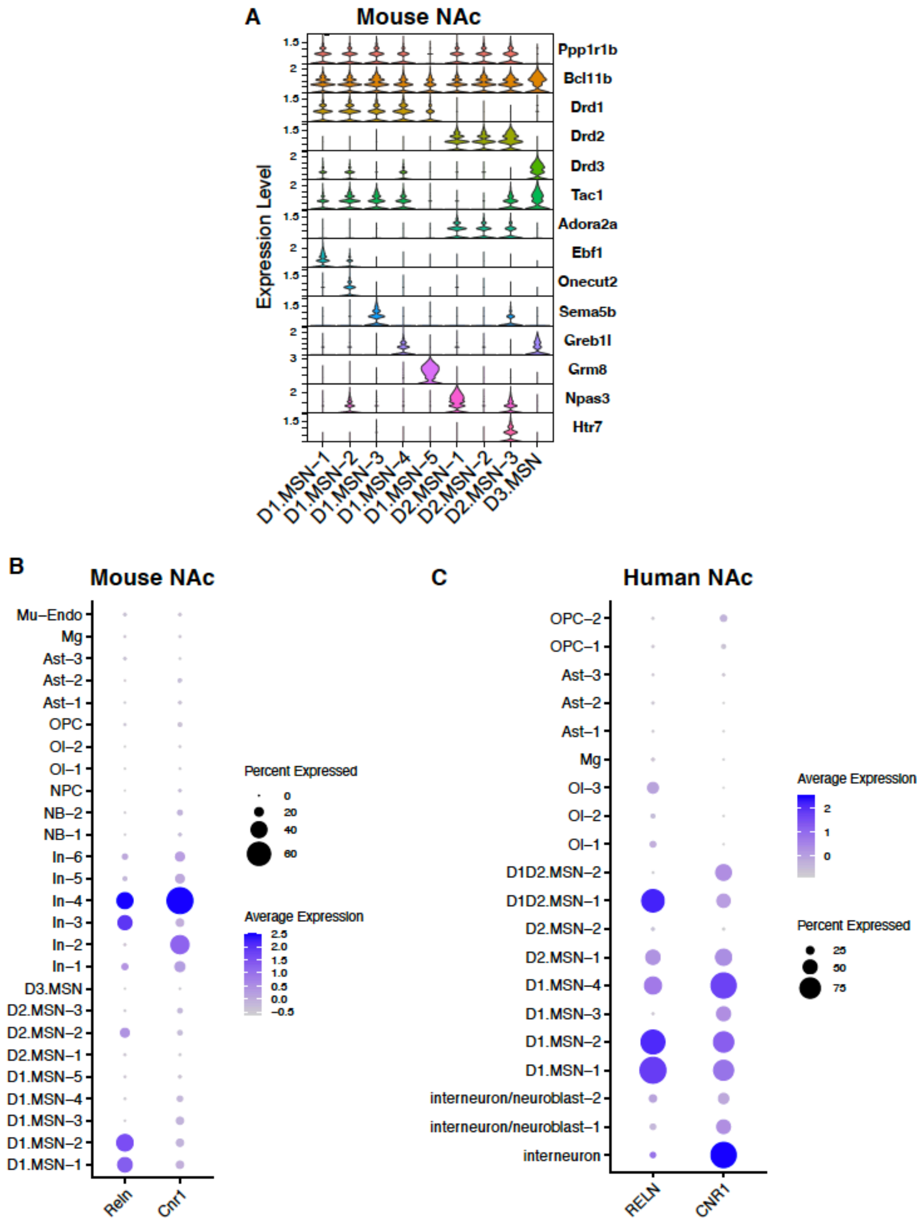


**Fig. S5. Medium spiny neuron subtype markers and cell type–specific *Reln/Cnr1* expression in the nucleus accumbens.** (**A**) Violin plots showing expression levels of selected marker genes across MSN subtypes in the adolescent mouse NAc, illustrating transcriptional distinctions among D1-, D2-, and D3-MSN populations. (**B**) Dot plot showing expression of *Reln* and *Cnr1* across mouse NAc cell types. Dot size represents the percentage of cells expressing each gene, and color intensity indicates average expression level. (**C**) Dot plot showing expression of RELN and CNR1 across cell types in the adult human NAc, displayed using the same scaling conventions as in (B), highlighting conserved cell type–specific expression patterns across species.

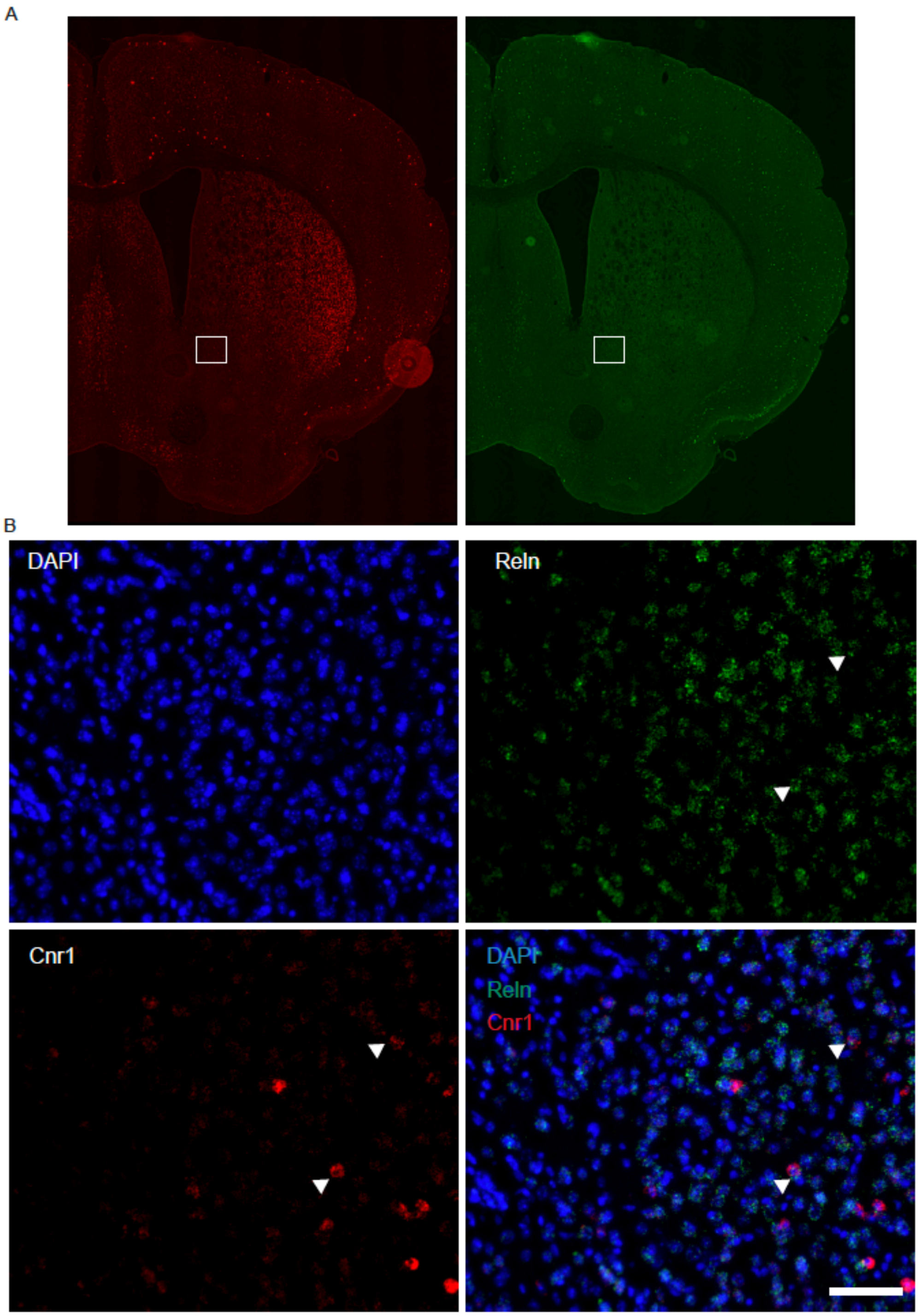


**Fig. S6. Spatial validation of *Reln* and *Cnr1* co-expression in mouse ventral striatum by multiplexed RNAscope.** (**A**) Low-magnification (2×) RNAscope images of a coronal mouse brain section showing expression of *Reln* (green) and *Cnr1* (red). The boxed region indicates the area shown at higher magnification below. (**B**) High-magnification views of the boxed region showing DAPI (blue), *Reln* (green), *Cnr1* (red), and merged channels. Arrowheads indicate cells exhibiting co-expression of *Reln* and *Cnr1*. Scale bar 100 µm

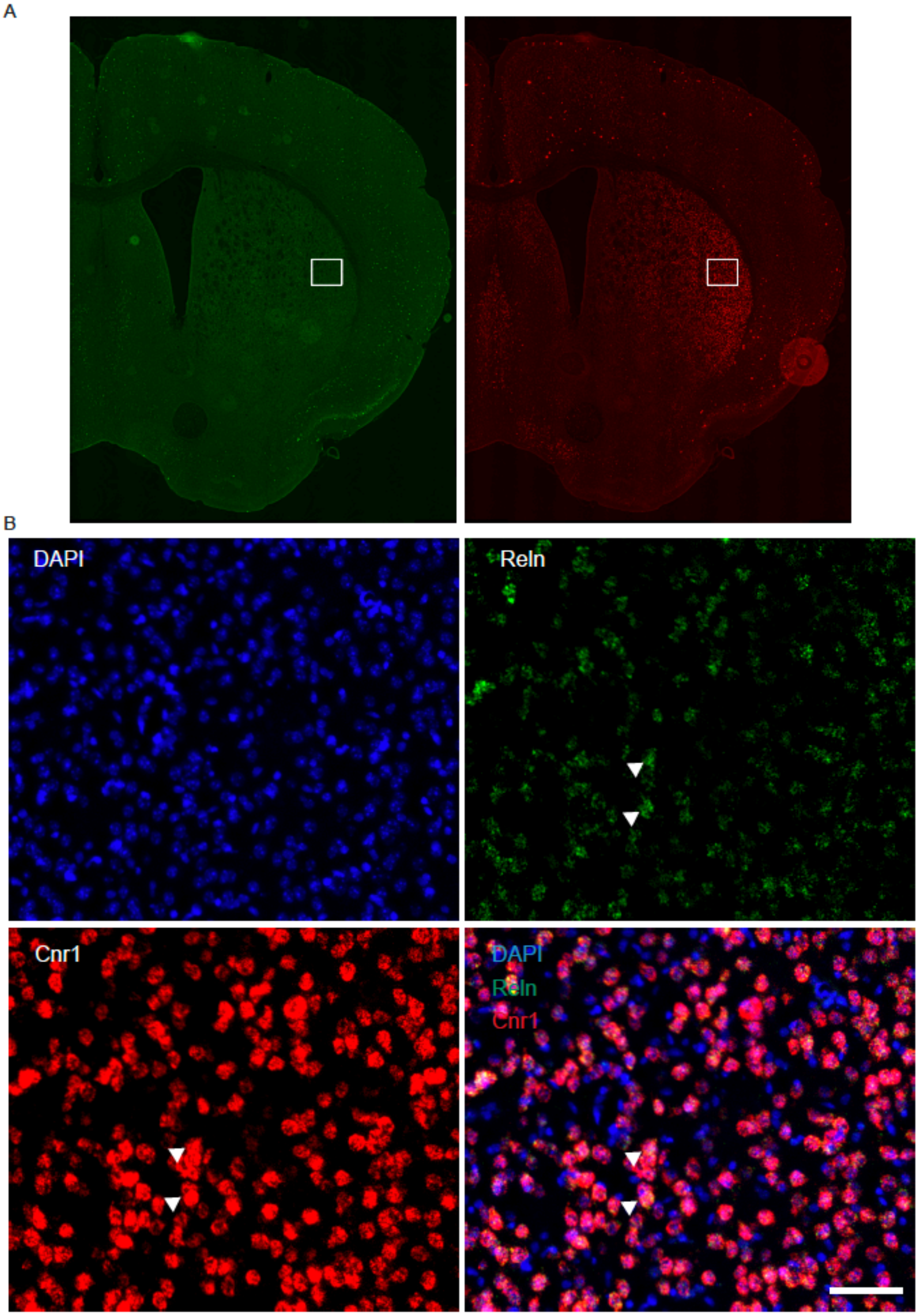


**Fig. S7. Spatial validation of *Reln* and *Cnr1* co-expression in mouse dorsolateral striatum by multiplexed RNAscope.** (**A**) Low-magnification (2×) RNAscope images of a coronal mouse brain section showing expression of *Reln* (green) and *Cnr1* (red). The boxed region indicates the area shown at higher magnification below. (**B**) High-magnification views of the boxed region showing DAPI (blue), *Reln* (green), *Cnr1* (red), and merged channels. Arrowheads indicate cells exhibiting co-expression of *Reln* and *Cnr1*. Scale bar 100 µm

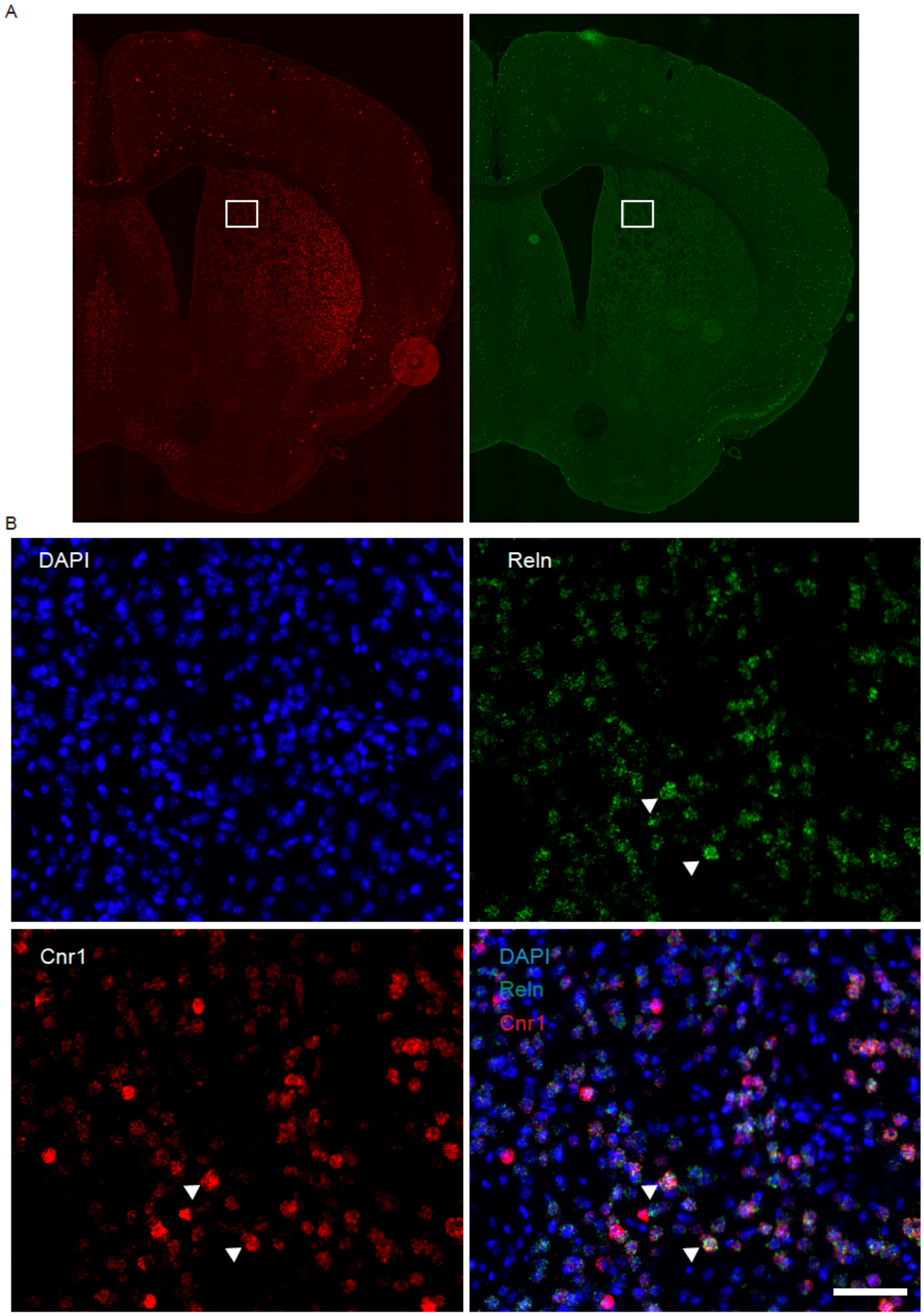


**Fig. S8. Spatial validation of *Reln* and *Cnr1* co-expression in mouse dorsomedial striatum by multiplexed RNAscope.** (**A**) Low-magnification (2×) RNAscope images of a coronal mouse brain section showing expression of *Reln* (green) and *Cnr1* (red). The boxed region indicates the area shown at higher magnification below. (**B**) High-magnification views of the boxed region showing DAPI (blue), *Reln* (green), *Cnr1* (red), and merged channels. Arrowheads indicate cells exhibiting co-expression of *Reln* and *Cnr1*. Scale bar 100 µm

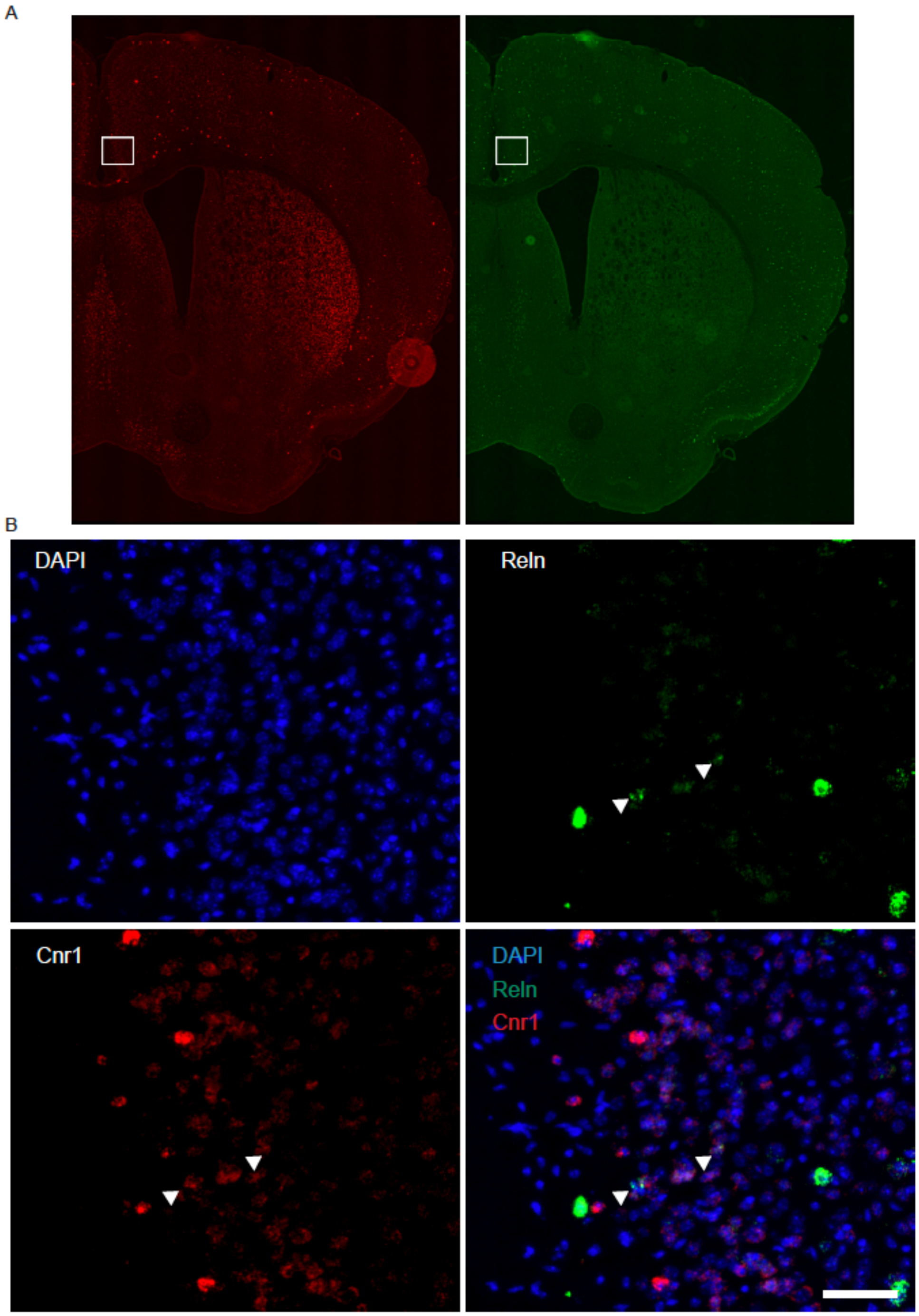


**Fig. S9. Spatial validation of *Reln* and *Cnr1* co-expression in mouse cortex by multiplexed RNAscope.** (**A**) Low-magnification (2×) RNAscope images of a coronal mouse brain section showing expression of *Reln* (green) and *Cnr1* (red). The boxed region indicates the area shown at higher magnification below. (**B**) High-magnification views of the boxed region showing DAPI (blue), *Reln* (green), *Cnr1* (red), and merged channels. Arrowheads indicate cells exhibiting co-expression of *Reln* and *Cnr1*. Scale bar 100 µm

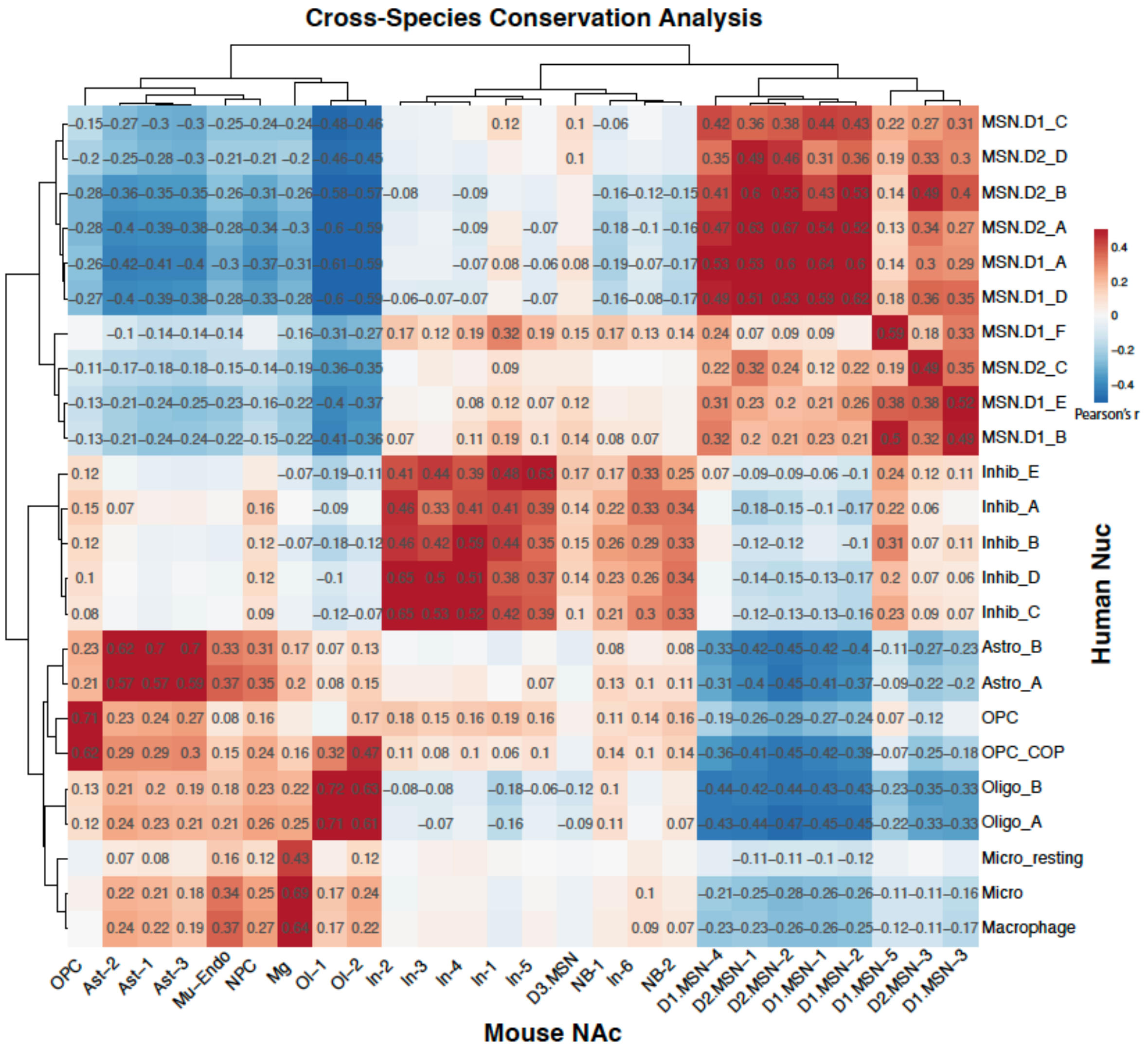


**Fig. S10. Cross-species conservation of nucleus accumbens cell types.** Heatmap showing Pearson correlation coefficients between adolescent mouse NAc cell classes (columns) and adult human NAc cell populations (rows). Correlations were computed using t-statistics derived from the top 100 marker genes for each cell type, restricted to genes with annotated mouse-human orthology. Hierarchical clustering highlights conserved transcriptional relationships across species. Color scale indicates Pearson's *r*.

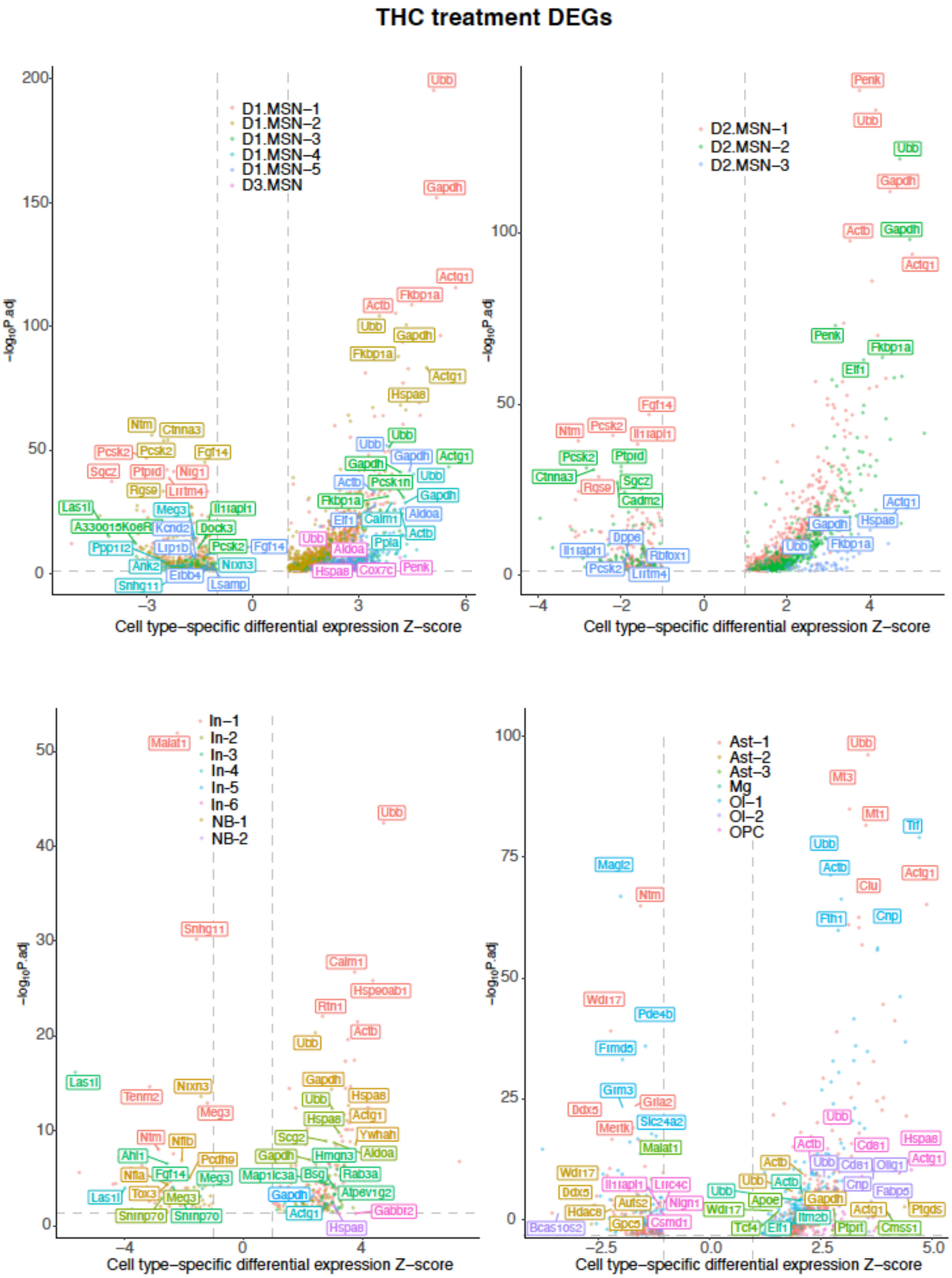


**Fig. S11. Cell type-specific transcriptional responses to adolescent THC exposure.** Volcano plots showing DEGs induced by THC exposure across major NAc cell classes, including D1-MSNs, D2-MSNs, interneurons (In), neuroblasts (NB), and non-neuronal cell types. The x-axis represents cell type-specific differential expression z-scores, and the y-axis shows -log$_{10}$(adjusted *p* value). Top significantly regulated genes are labeled.

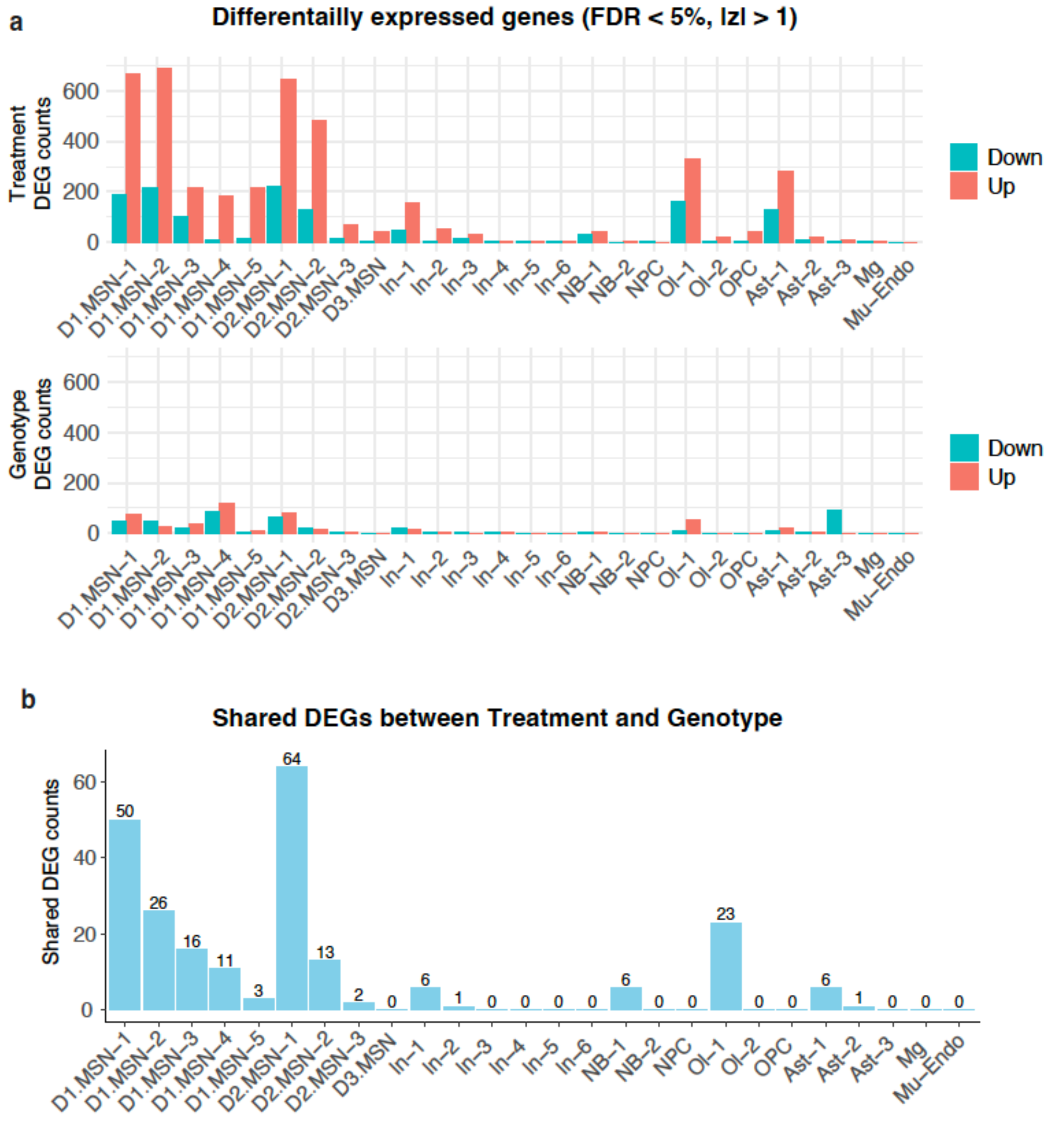


**Fig. S13. Shared and distinct transcriptional responses to THC exposure and *Reln* haploinsufficiency across NAc cell types.** (**A**) Bar plots showing the number of differentially expressed genes (DEGs; FDR < 5%, |z| > 1) identified per cell type following adolescent THC exposure (top) or *Reln* haploinsufficiency (bottom). Upregulated and downregulated genes are shown separately. (**B**) Number of DEGs shared between THC treatment and *Reln* haploinsufficiency within each cell type.

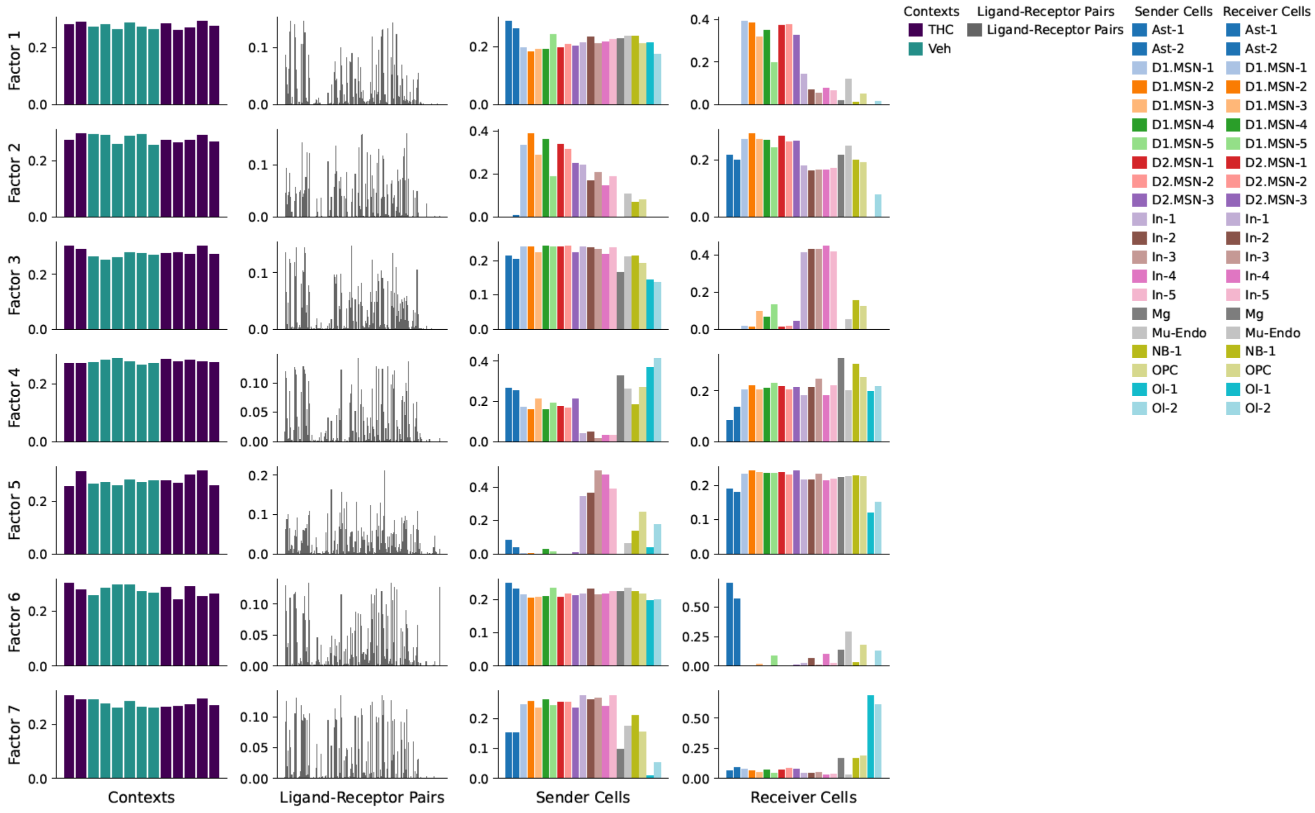


**Fig. S14. Tensor-based decomposition of cell-cell communication programs in the adolescent NAc following THC exposure.** Cell-cell communication was inferred using LIANA and decomposed using tensor component analysis (Tensor-cell2cell), yielding seven latent factors (rows), each representing a distinct intercellular communication program. For each factor, loadings (y-axis) are shown for each tensor dimension (columns), including experimental context (THC vs vehicle), ligand-receptor pairs, sender cell types, and receiver cell types (x-axis). Bars are colored by major cell-type classes as indicated. Differences in context loadings highlight communication programs that are selectively associated with THC exposure relative to vehicle controls.

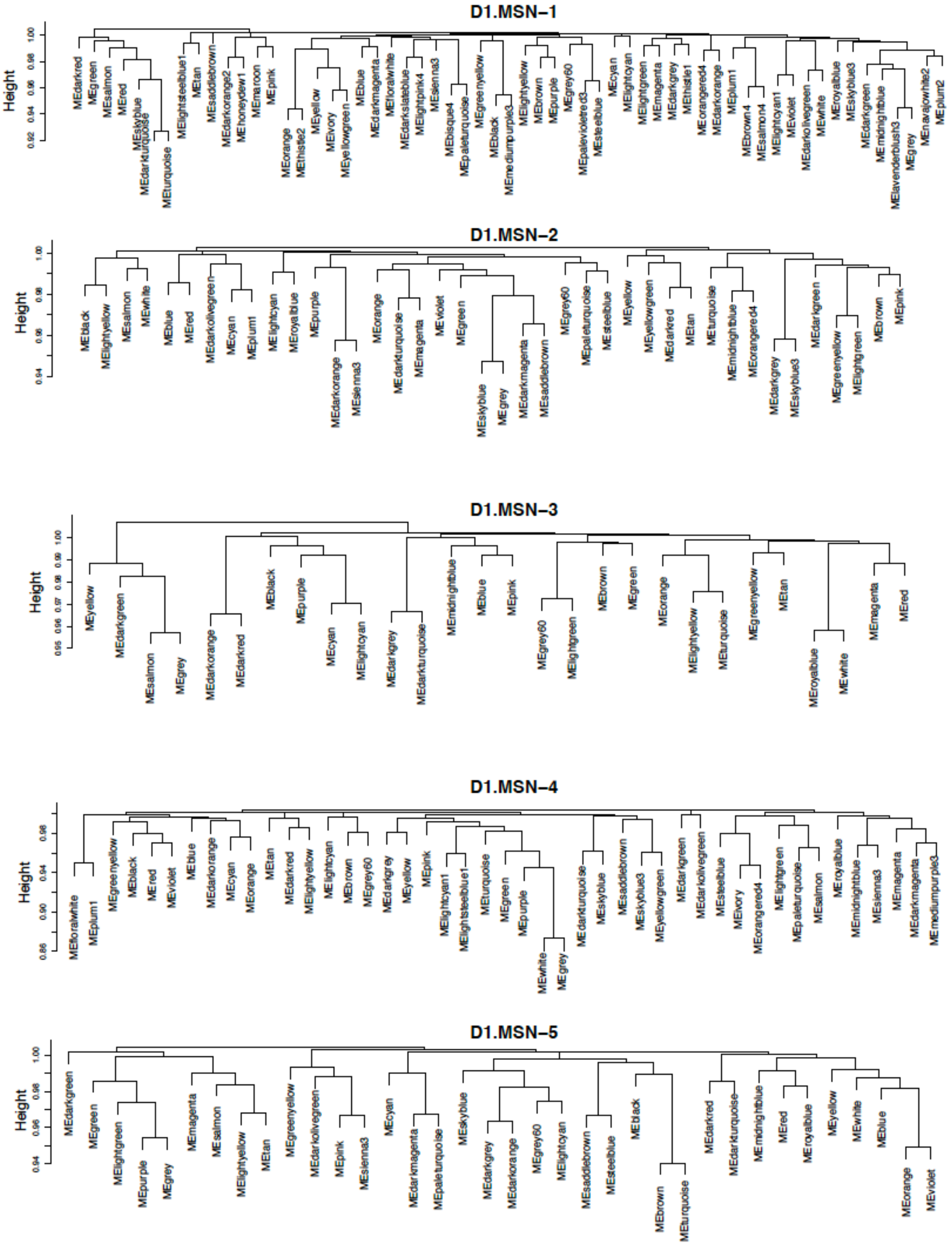


**Fig. S15. Hierarchical clustering of gene co-expression modules in D1 medium spiny neuron subtypes.** Dendrograms show hierarchical clustering of module eigengenes identified by WGCNA within each D1-MSN subtype (D1.MSN-1 through D1.MSN-5). Branch height reflects eigengene dissimilarity, illustrating relationships among transcriptional modules within each MSN subtype.

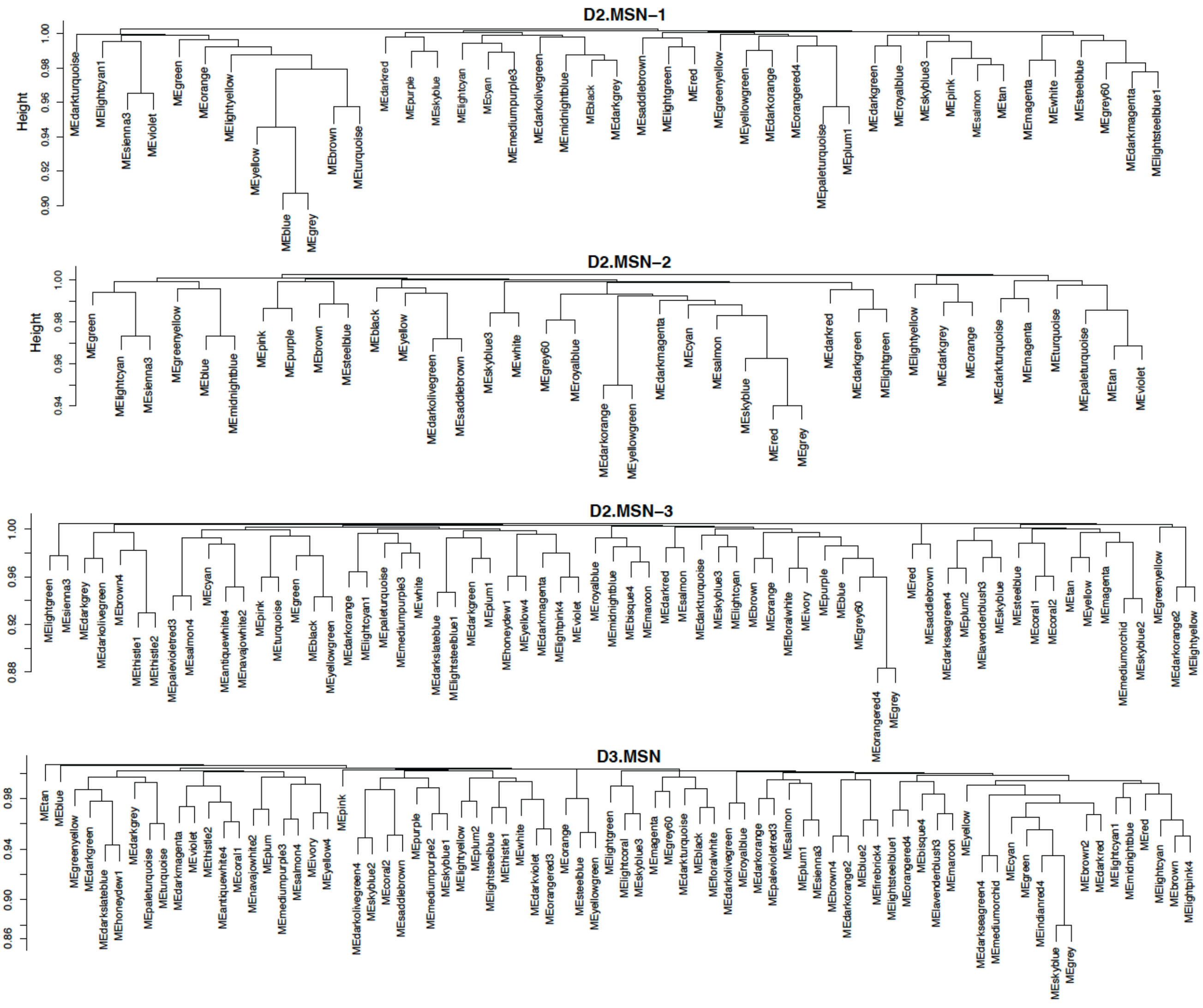


**Fig. S16. Hierarchical clustering of gene co-expression modules in D2 medium spiny neuron subtypes.** Dendrograms show hierarchical clustering of module eigengenes identified by WGCNA within each D2-MSN subtype (D2.MSN-1, D2.MSN-2, D2.MSN-3) and D3.MSNs. Branch height reflects eigengene dissimilarity, illustrating relationships among transcriptional modules within each MSN subtype.

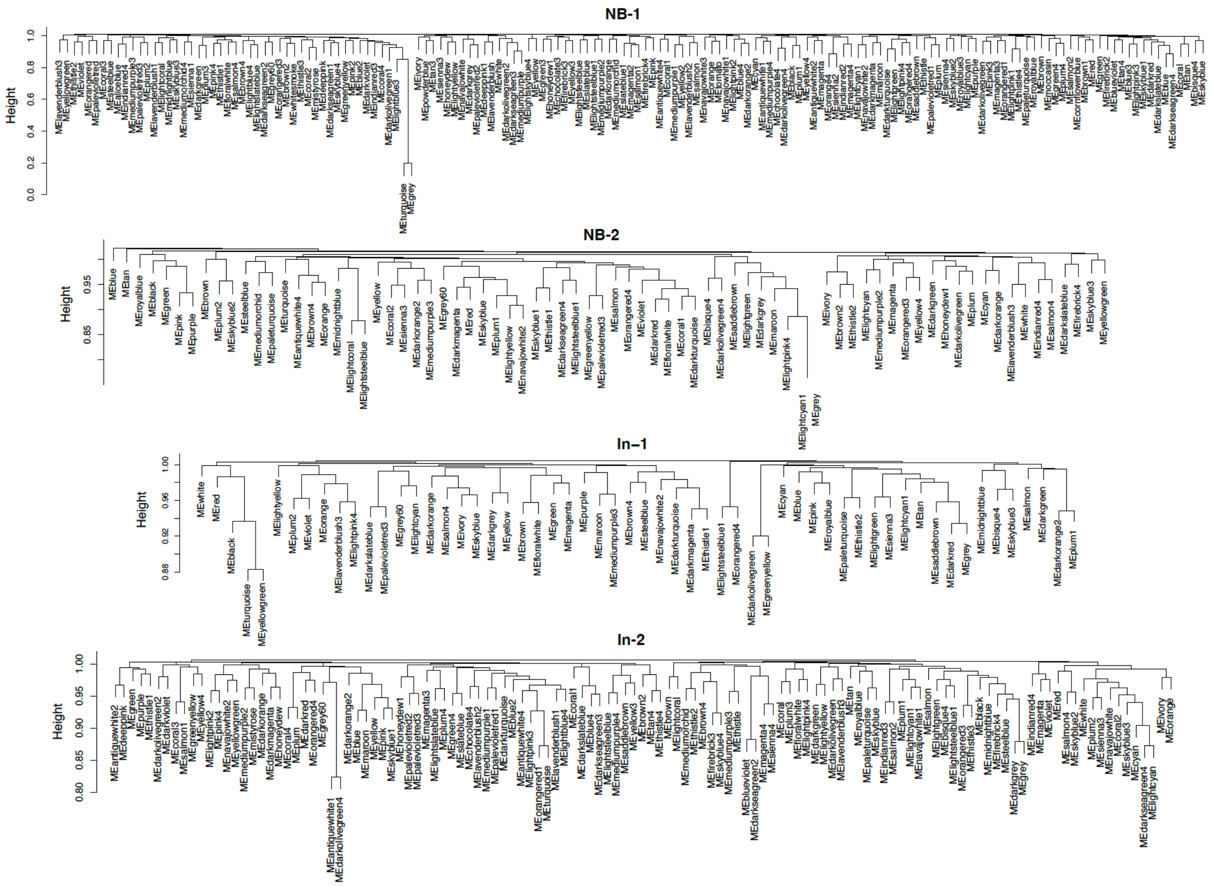


**Fig. S17. Hierarchical clustering of gene co-expression modules in neuroblast and interneuron populations.** Hierarchical clustering dendrograms of module eigengenes identified by WGCNA within neuroblast subtypes (NB-1, NB-2) and interneuron subtypes (In-1, In-2). Dendrogram branch height reflects eigengene dissimilarity, illustrating relationships among transcriptional modules within each cell population.

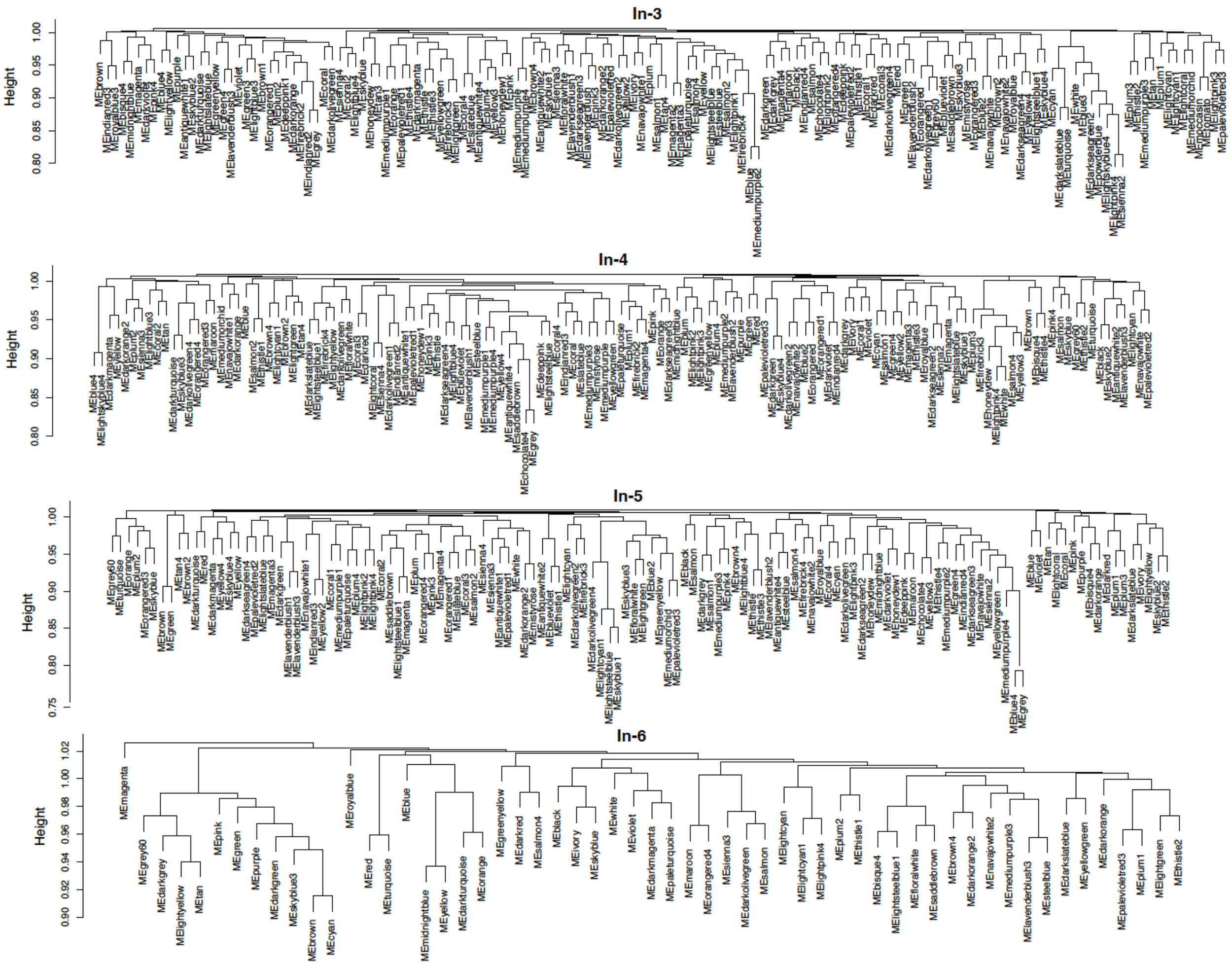


**Fig. S18. Hierarchical clustering of gene co-expression modules in interneuron subtypes.** Hierarchical clustering dendrograms of module eigengenes derived from WGCNA within interneuron subtypes In-3, In-4, In-5, and In-6. Dendrogram height reflects eigengene dissimilarity, illustrating relationships among transcriptional modules within each interneuron population.

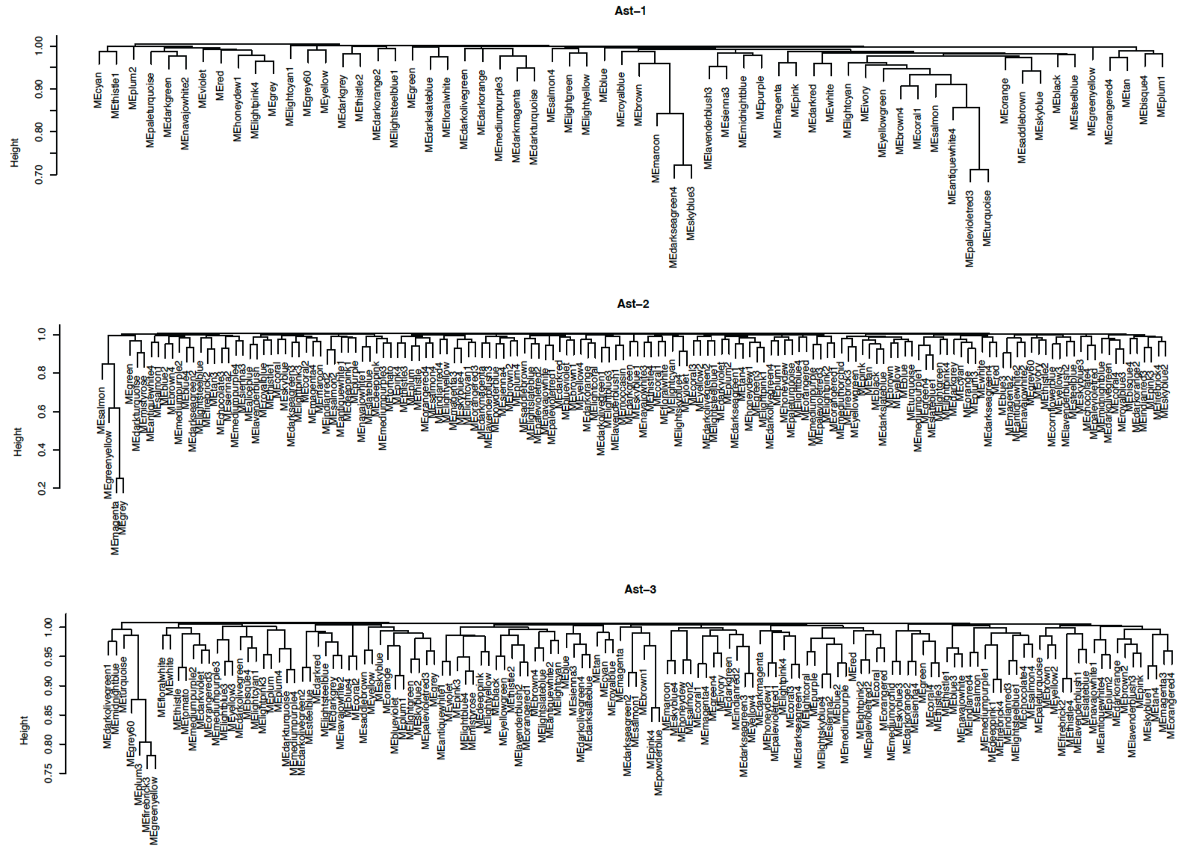


**Fig. S19. Hierarchical clustering of module eigengenes in astrocyte subtypes.** Hierarchical clustering dendrograms of WGCNA module eigengenes for astrocyte subtypes Ast-1, Ast-2, and Ast-3. Dendrograms depict the similarity structure among module eigengenes within each astrocyte population based on pairwise correlations, with branch height indicating eigengene dissimilarity.

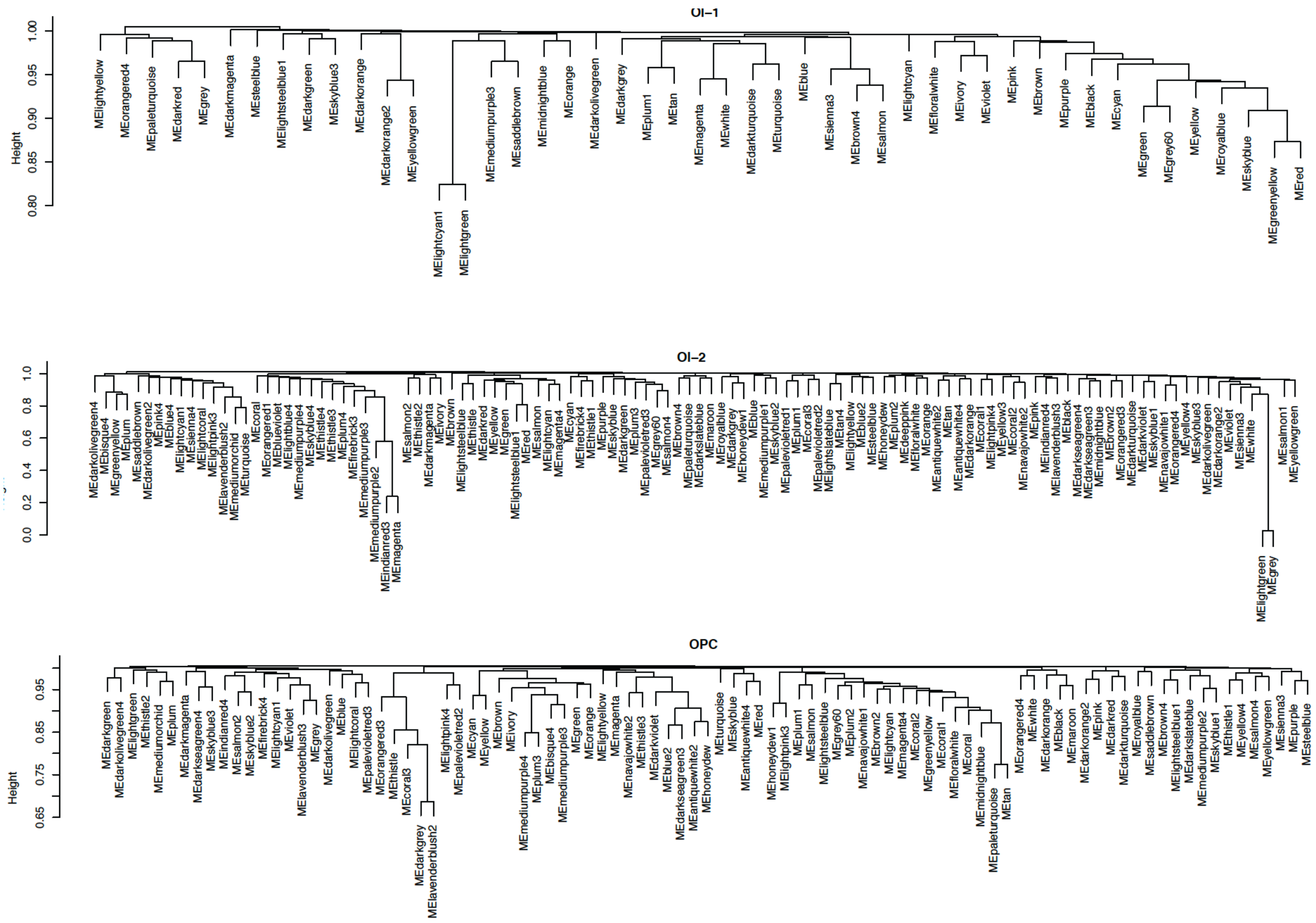


**Fig. S20. Hierarchical clustering of module eigengenes in oligodendrocyte subtypes.** Hierarchical clustering dendrograms of WGCNA module eigengenes for oligodendrocyte subtypes Ol-1, Ol-2, and OPC. Dendrograms depict the similarity structure among module eigengenes within each oligodendrocyte population based on pairwise correlations, with branch height indicating eigengene dissimilarity.

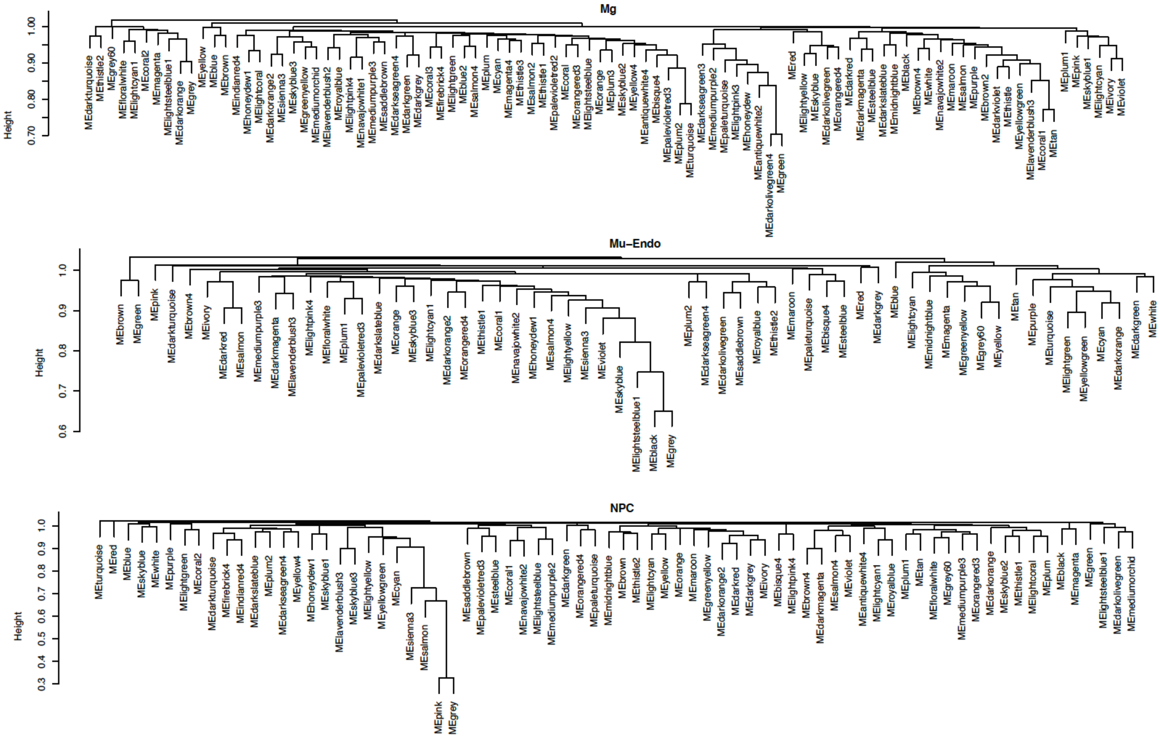

**Fig. S21. Hierarchical clustering of module eigengenes in microglia, mu-endo, NPC subtypes.** Hierarchical clustering dendrograms of WGCNA module eigengenes for other subtypes, including microglia, mu-endo, NPC subtypes. Dendrograms depict the similarity structure among module eigengenes within each population based on pairwise correlations, with branch height indicating eigengene dissimilarity.

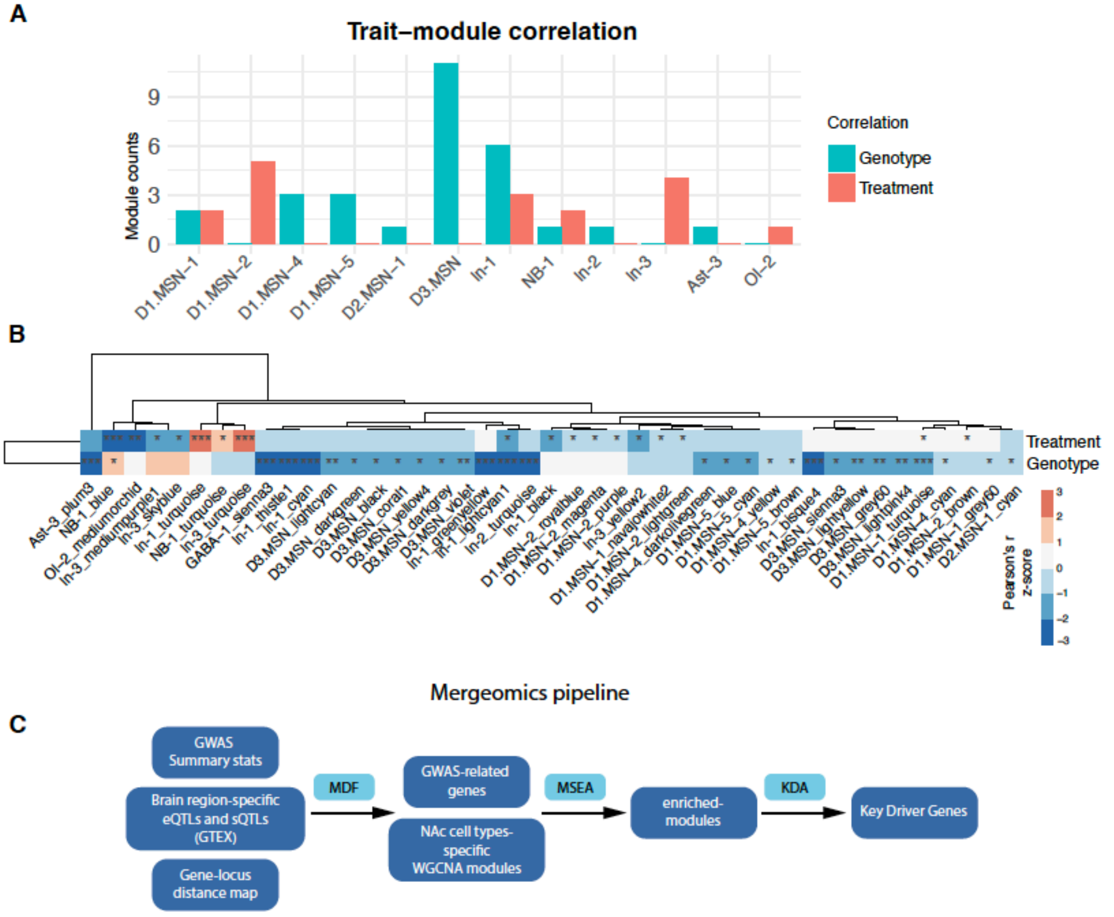


**Fig. S22. Trait-module associations and integration with human genetic risk.** (**A**) Number of gene co-expression modules showing significant correlation with genotype or treatment across cell types. (**B**) Heatmap of module–trait correlations between WGCNA-derived gene modules and experimental conditions (treatment and genotype). Color indicates Pearson's correlation coefficient; significance is denoted as *p < 0.05, **p < 0.01, ***p < 0.001. (**C**) Schematic of the Mergeomics analytical pipeline used to integrate NAc cell type-specific gene modules with human GWAS data. GWAS summary statistics and brain region-specific eQTL/sQTL annotations (GTEx) were combined with gene-locus distance maps and WGCNA modules, followed by marker dependency filtering (MDF), marker set enrichment analysis (MSEA), and key driver analysis (KDA) to identify enriched modules and candidate key driver genes.

**Data S1. Sample annotation and sequencing summary for snRNA-seq libraries. (separate file)**
Table listing sample identifiers, sex, genotype, treatment condition, estimated number of nuclei recovered, mean reads per nucleus, and median genes detected per nucleus for all single-nucleus RNA-seq libraries analyzed in this study.

**Data S2. Cell-level metadata for single-nucleus RNA-seq analysis. (separate file)**
Table containing per-nucleus quality control metrics, sample identity, genotype and treatment annotations, mitochondrial and ribosomal content, doublet classification, SCTransform-normalized values, clustering assignments, and final detailed and coarse cell-type annotations for all nuclei included in the analysis.

**Data S3. Differential gene expression analysis by cell population for genotype. (separate file)**
Each worksheet corresponds to a single cell type and reports differentially expressed genes identified by *FindMarkers* for *Reln* genotype effects.

**Data S4. Differential gene expression analysis by cell population for treatment. (separate file)**
Each worksheet corresponds to a single cell type and reports differentially expressed genes identified by *FindMarkers* for THC treatment effects.

**Data S5. Overlap between Factor 3-associated genes and DEGs. (separate file)**
Table listing the 46 genes shared between the top 100 genes with highest Factor 3 loadings from tensor-based cell-cell communication analysis and the set of treatment DEGs.

**Data S6. Cell type-specific WGCNA module gene lists. (separate file)**
Each worksheet contains the genes comprising individual WGCNA modules identified within each annotated cell population.

**Data S7. Cell type-specific trait-module associations. (separate file)**
The worksheet reports Pearson correlations between module eigengenes, and experimental variables derived from WGCNA for each cell type.
